# A deep genetic analysis of banana Fusarium wilt pathogens of Cuba in a Latin American and Caribbean diversity landscape

**DOI:** 10.1101/2023.08.29.553192

**Authors:** Einar Martínez de la Parte, Luis Pérez-Vicente, David E. Torres, Anouk van Westerhoven, Harold J. G. Meijer, Michael F. Seidl, Gert H. J. Kema

**Affiliations:** Laboratory of Phytopathology, Wageningen University & Research, The Netherlands; Instituto de Investigaciones de Sanidad Vegetal (INISAV), Ministry of Agriculture, Cuba; Theoretical Biology and Bioinformatics Group, Department of Biology, Utrecht University, The Netherlands; BU Biointeractions and Plant Health, Wageningen Plant Research, Wageningen University & Research, Wageningen, The Netherlands

**Keywords:** Banana, Fusarium, phylogeography, genotyping, pathogenicity, races

## Abstract

- Fusarium wilt of bananas (FWB) is a devastating plant disease that causes significant economic losses in banana production worldwide and is one of the major concerns for Cuban banana cultivation. The disease is caused by members of the soil-borne *Fusarium oxysporum* species complex. However, the genetic diversity among *Fusarium* species infecting bananas in Cuba is currently unknown.
- We conducted a comprehensive survey of symptomatic banana plants across all production zones of the country and assembled a collection of 170 *Fusarium* isolates. Using genotyping-by-sequencing and whole-genome comparisons, we investigated the genetic diversity across this suite of isolates and compared it with the genetic diversity of a global *Fusarium* panel.
- Typical FWB symptoms were observed in varieties of the Bluggoe cooking banana and Pisang Awak subgroups in 14 provinces. Phylogenetic analysis revealed that *F. purpurascens, F. phialophorum,* and *F. tardichlamydosporum* cause FWB in Cuba, with the latter dominating the population. Furthermore, we identified between five and seven genetic clusters, with *F. tardichlamydosporum* isolates divided into at least two distinct subgroups, indicating a high genetic diversity of *Fusarium* spp. causing FWB in the Americas.
- Our study provides unprecedented insights into the population genetic structure and diversity of the FWB pathogen in Cuba and the Latin American and Caribbean regions.

## INTRODUCTION

Bananas are among the most produced, traded, and consumed fruits globally (FAO, 2022). In terms of global production, bananas directly follow behind major crops such as wheat, rice and maize (Perrier *et al*., 2011). With more than 1,000 cultivars, bananas are a staple food for more than 400 million people who rely on the commodity for food security and income. Approximately 84% of the crop is grown by smallholders and delivered to domestic markets, while the remaining 16% goes to international markets with an estimated annual export volume of approximately 20 million tons (FAO, 2022).

The global production of bananas, including plantains, is threatened by a range of different globally spreading plant diseases (Drenth & Kema, 2021). Among these, Fusarium wilt of banana (FWB) is one of the most destructive and widespread diseases of the crop (Ploetz, 2015a). It wiped out the famed Gros Michel variety in the previous century and currently devastates Cavendish plantations, and many other banana cultivars consumed and traded locally around the world (Ploetz, 2019; Staver *et al*., 2020; Westerhoven *et al*., 2022). The disease is caused by diverse, soil-borne anamorphic fungi belonging to the *Fusarium oxysporum* species complex (FOSC), which have been traditionally classified as *Fusarium oxysporum* f. sp. *cubense* (*Foc*). Genetic diversity was previously described by using vegetative compatibility groups (VCGs; Ploetz & Correll, 1988; Brake *et al*., 1990; Ploetz, 1990), a system where genetically related individuals exchange genetic material through heterokaryosis (Pérez-Vicente *et al*., 2014). Strains with identical alleles at vegetative or heterokaryon incompatibility (*het*/*vic*) loci belong to the same VCG (Correll, 1991). Physiological specialization towards particular banana varieties resulted in the definition of a core set of races; Race 1, Race 2 and Race 4, the latter being divided into Subtropical Race 4 (STR4) and Tropical Race 4 (TR4). Race 1 strains have been identified in no less than nine VCGs, are avirulent to Cavendish under regular conditions, but can cause FWB under abiotic stress and are then categorized as STR4 (Buddenhagen, 2009; Ploetz, 2015b; Mostert *et al*., 2017). This illustrates that there is no strict correlation between VCGs and races in the FWB-banana pathosystem (Ordóñez *et al*., 2015). Nevertheless, VCGs (up to 24) have been used to gain insight into the dissemination and evolution of the causal agents of FWB (Fourie *et al*., 2009; Ordóñez *et al*., 2015; Mostert *et al*., 2022), although they do not provide a measure of genetic distance between phenotypes (Bentley *et al*., 1998; Fourie *et al*., 2009). Thus, neither VCGs nor physiological races have contributed to a thorough understanding of the phylogeny and dissemination of the causal fungi of FWB (Dita *et al*., 2020).

DNA-based techniques provide detailed information about the genetic distance of different pathogen populations (Freese & Beyhan, 2023; Feurtey *et al*., 2023). Previously, phylogenetic analyses revealed that *Foc* is polyphyletic and divided over the three recognized FOSC clades, which can be further divided into eight to 10 individual genetic lineages (Bentley *et al*., 1998; O’Donnell *et al*., 1998; Groenewald *et al*., 2006; Fourie *et al*., 2009; Mostert *et al*., 2017; Ordoñez, 2018). High-resolution genotyping-by-sequencing analyses using Diversity Array Technology (DArTseq; Cruz *et al*., 2013, Kilian *et al*., 2003) is a genome-wide method that validated and extended these findings (Ordóñez *et al*., 2015; Mostert *et al*., 2022) It has also been used to study genetic diversity and resistance mapping in many crop species, such as cassava, chickpea, pea, tea, and wheat (Sohail *et al*., 2015; Malebe *et al*., 2019; Adu *et al*., 2021; Sampaio *et al*., 2021; Ahmed *et al*., 2021) as well as in banana (Amorim *et al*., 2009; Risterucci *et al*., 2009; Sardos *et al*., 2016; Ahmad *et al*., 2020) and their pathogens (Sharma *et al*., 2014; Ordóñez *et al*., 2015; Mostert *et al*., 2022).

Bananas arrived in Cuba in the 16^th^ century and are among the most produced fruits in the country, required for food security and providing an important source of income for small growers (ONEI, 2022; Martínez-de la Parte *et al*., 2023). Fusarium wilt of banana has been present in Cuba since the end of 19^th^ century (Battle & Pérez-Vicente, 2009) and contributed significantly to the collapse of the exporting activities during the first-mid of the last century. After the substitution of the susceptible ‘Gros Michel’ (AAA) by resistant Cavendish (AAA) varieties during the 1950s and the large-scale cultivation of plantain (AAB) varieties, FWB lost its economical relevance (Battle & Pérez-Vicente, 2009). However, in smallholder farms and backyard gardens with conducible soils, planted with susceptible cultivars, the disease is still present. In a previous survey, FWB was observed in eight provinces across a suite of important banana cultivars in Cuba, which resulted in a small collection of 52 isolates that was characterized and divided into four VCGs and two races (Battle & Pérez-Vicente, 2009). Recently, the devastating TR4 was reported in Colombia, Venezuela, and Peru (Garcia-Bastidas *et al*., 2020; Acuña *et al*., 2021; Herrera *et al*., 2023) and threatens Cuban production as 56% of the cultivated germplasm is highly susceptible to this race (Martínez de la Parte *et al*., 2023). We, therefore, initiated a nationwide survey of many geographically and environmentally different locations, resulting in a collection of 170 isolates that was subsequently characterized by DArTseq and whole-genome sequencing. These analyses enabled us to decipher the FWB causing Fusarium species in Cuba, and to place local diversity into the greater diversity landscape across Latin America and the Caribbean (LAC).

## MATERIALS AND METHODS

### Survey and sampling

A comprehensive survey of FWB was undertaken in Cuba between 2016-2018. In total, locations in 58 municipalities of 15 provinces were visited, representing the main banana-producing regions in Cuba. Plants with FWB symptoms - including chlorotic and wilted leaves, collapsed leaves at the petioles, pseudostem discoloration, and splitting - were sampled from large banana plantations, smallholder farms, and backyard gardens (Figure 1). At each site, global positioning coordinates and altitude data were collected, and the varieties were identified. The pseudostems of the diseased plants were cut and discolored vascular strands were collected, placed on sterile filter paper to dry, and packed in a paper envelope until further analyses.

**Fig. 1.**
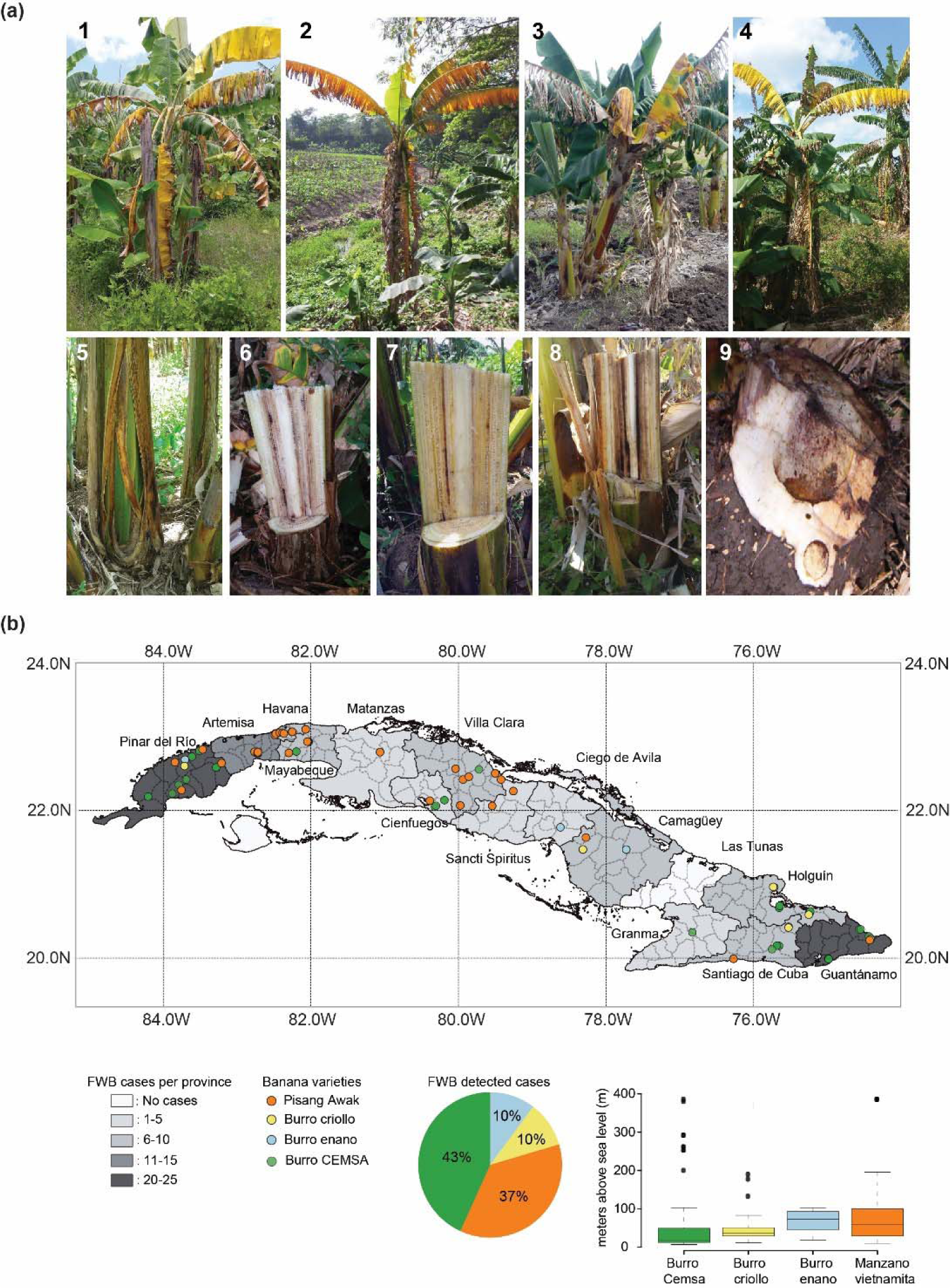
Fusarium wilt of banana (FWB) is widely distributed across Cuba. (A) Typical symptoms of FWB detected during the survey; leaf chlorosis or collapse in banana cultivars ‘Pisang Awak’ [1], ‘Burro Criollo’ [2], ‘Burro Enano’ [3], and in ‘Burro CEMSA’ [4]. Pseudostem splitting in Burro CEMSA [5], internal pseudostem symptoms (discolored vascular strands) in ‘Pisang Awak’ [6] and in ‘Burro CEMSA’ [7, 8]. Corm necrosis in ‘Burro Criollo’ [9]. (B) Overview of the 2016-2018 survey. The FWB incidence is indicated by the shading of the provinces and colored dots represent different banana varieties, orange dots are ‘Pisang Awak’ fields with FWB symptoms, yellow dots are fields with FWB affected ‘Burro Criollo’, blue dots are locations with FWB-affected ‘Burro Enano’ plants, and green dots represent FWB cases in ‘Burro CEMSA’. The pie chart indicates the proportion of FWB observed in four banana varieties that are cultivated in Cuba, which collectively represent 44.4% on the national banana acreage.

### Strain isolation, purification, and maintenance

Dried pseudostem samples were cut into pieces of ∼0.8 cm^2^, surface sterilized with ethanol (70%) and commercial bleach (1%) for 1 min each, rinsed with sterile distilled water, plated on 2% water agar (WA) supplemented with streptomycin sulphate (100 µg/mL) in 9 cm-diameter Petri dishes, and incubated at 25°C in complete darkness. After approximately two days, fungal colonies resembling *Fusarium* were transferred to potato dextrose agar plates (PDA, 24g/L; Thermo Scientific™ Oxoid™, Landsmeer, the Netherlands). A total of 40 isolates from a previous survey in 1993-1994 (Battle-Viera and Pérez-Vicente, 2009) were also included in the collection, resulting in a Cuban panel of 170 isolates (Table S1).

Single-spore colonies were generated for each isolate and these were maintained in PDA for short-term use and as seven-day-old spore suspensions in glycerol stocks (2mL, 15%) for long-term preservation at −80°C in Nunc® CryoTubes® (Sigma-Aldrich, Roskilde, Denmark). The entire collection was deposited in the microbial culture collection of the Instituto de Investigaciones de Sanidad Vegetal (INISAV, Cuba) with a copy at the Laboratory of Phytopathology of Wageningen University & Research (WUR; Table S1). The total data set that was analyzed in this study comprised the Cuban panel of 170 isolates, complemented by a collection of 210 isolates from the Americas and a set of 33 isolates representing a global panel that was analyzed earlier (Ordóñez *et al*., 2023).

### DNA isolation and PCR amplifications

Fungal isolates were grown on PDA plates for seven days at 28°C. Then, conidiospores were collected and transferred to a flask with 100 mL potato dextrose broth (PDB, 34g/L) (Thermo Scientific™ Oxoid™, Landsmeer, the Netherlands, Landsmeer, the Netherlands) and incubated by shaking at 140 rpm and 25°C for 5 days. The mycelium was obtained by filtering the inoculum through two layers of sterile Miracloth and washed at least twice with sterile MQ water. Subsequently, the mycelium was freeze-dried overnight in a 2 mL Eppendorf tube and ground in a mortar with a pestle using liquid nitrogen. Genomic DNA of each isolate was extracted using the MasterPure™ Yeast DNA Purification Kit (LGC Biosearch Technologies, Halle-Zoersei, Belgium), according to the manufactureŕs protocol. Genomic DNA size, concentration and integrity were assessed by gel electrophoresis, NanoDrop™ One/OneC Microvolume UV-Vis Spectrophotometer (Thermo Scientific™, Landsmeer, the Netherlands) and Qubit analysis (Qubit™ Flex Fluorometer, Thermo Fisher Scientific, Landsmeer, the Netherlands).

Primer pairs PFO2 (5’-CCCAGGGTATTACACGGT-3’) and PFO3 (5’-CGGGGGATAAAGGCGG-3’) (Edel *et al*., 2000), FocTR4-F (CACGTTTAAGGTGCCATGAGAG) and Foc TR4-R (5’-GCACGCCAGGACTGCCTCGTGA-3’) (Dita *et al*., 2010), Six1a_266-F (5’-GTGACCAGAACTTGCCCACA-3’) and Six1a_266-2R (5’-CTTTGATAAGCACCATCAA-3’) and Six6b_210-F (5’-ACGCTTCCCAATACCGTCTGT-3’) and Six6b_210-R (5’-AAGTTGGTGAGTATCAATGC-3’) (Carvalhais *et al*., 2019) were used to identify the isolates, using the described conditions for PCR in 25 µL using GoTaq® G2 DNA Polymerase and Master Mix (Promega Benelux BV, Leiden, the Netherlands), 10 µM of each primer, 10–20 ng of DNA, and sterile deionized water. The amplicons were resolved by electrophoresis using 1.5% agarose gel, stained with ethidium bromide, visualized and photographed using the Chemidoc^TM^ MP image system (Bio-Rad Laboratories, Veenendaal, the Netherlands). Amplicon sizes were estimated using a 100-bp ladder as reference.

Mating type idiomorphs were determined by PCR using the primer set Falpha 1 (5’-CGGTCAYGAGTATCTTCCTG-3’) and Falpha 2 (5’-GATGTAGATGGAGGGTTCAA-3’) for *mat1-1* (Arie *et al*., 2000), and FF1(5’-GTATCTTCTGTCCACCACAG-3’) and Gfmat2c (5’-AGCGTCATTATTCGATCAAG-3’) for *mat1-2* (Fourie *et al*., 2009). The cycling conditions for *MAT1-1* amplification were: initial denaturation at 95°C for 15 min; 35 cycles at 94°C for 1 min., 55°C for 30 s, 72°C for 1 min. and a final extension at 72°C for 10 min. The cycling conditions for *MAT1-2* were an initial denaturation at 95°C for 2 min; 35 cycles at 94°C for 1 min., 54°C for 40 s, 72°C for 2 min., and a final extension at 72°C for 7 min.

### Pathogenicity tests

Eleven strains representing different provinces and banana cultivars were tested for their pathogenicity towards Grand Naine (Cavendish-AAA, ITC0180), Gros Michel (Gros Michel-AAA, ITC1122), and Burro Cemsa (Bluggoe-ABB, ITC1259). Traditionally, race designations are based on the pathogenicity towards the aforementioned banana cultivars, where isolates causing FWB on ‘Gros Michel’ are considered as Race 1, on ‘Burro CEMSA’ as Race 2, and on ‘Grand Naine’ as TR4. Tissue culture plants, approximately 2.5-month-old, were inoculated with the selected strains following a previously published protocol (García-Bastidas *et al*., 2019). The *F. odoratissimum* reference strain II-5 (NRRL 54006, VCG01213) was included as a positive control, and plants treated with water were used as negative controls. All the assays were conducted in an environmentally controlled greenhouse compartment (28±2°C, 16h light, and ∼85% relative humidity) at the Unifarm greenhouse facility of WUR. Twelve weeks after inoculation, the corms of the plants were cut transversally, photographed and the Rhizome Discolored Area (RDA) was calculated using ImageJ 1.52r software (National Institutes of Health, Bethesda, MD, USA). At the end of the phenotyping experiments, a random banana plant per treatment was selected for re-isolation and (molecular) diagnosis of the FWB causal agent to complete Koch’s postulates.

### Diversity Array Technology by sequencing (DArTseq) analyses

DArTseq is a genome complexity reduction method by establishing genomic markers, which are scored as binary data to represent either the presence or absence of the marker sequence in each isolate (Jaccoud, 2001). Based on the DArTseq markers, single nucleotide polymorphisms (SNP) can be called and similarly encoded as binary data in a matrix. Isolated DNAs from monosporic cultures of each isolate of the Cuban panel were sent to Diversity Arrays Technology Pty. Ltd. (http://www.diversityarrays.com), Canberra, Australia, for genome complexity reduction and Illumina sequencing (Kilian *et al*., 2003), which was previously adapted for FOSC (Ordóñez *et al*., 2015). Additionally, we included markers that were generated in a collection of 243 strains from a global *Fusarium* diversity panel (Table S3), which represents the genetically distinct *Fusarium* species identified by Maryani *et al*. (2019).

The DArTseq markers and SNPs were analyzed using the R package dartR v.1.8.3 (Gruber *et al*., 2018; R Core Team, 2021), where ploidy was set as haploid, monomorphic loci were removed from both data sets, and only isolates with a repeatability index above 1 were maintained. Both the remaining loci and isolates were kept for downstream analyses when their call rates were higher than 0.95 (0.98 for SNPs) and >=10 positive markers or a MAF>=0.05 in the case of SNPs were present. Based on the filtered binary presence/absence matrix we constructed a distance matrix using the Dice similarity coefficient and inferred the phylogenetic relationships using the Neighbor Joining algorithm (bootstrap=1,000) and the previously generated distance matrix in R (R Core Team, 2021). Additionally, we performed a principal component analysis (PCA) with the filtered matrix to verify our results using factoextra and FactoMineR (Lê *et al*., 2008). To calculate the genetic diversity within each species in our dataset, we calculated the Bray-Curtis and Jaccard distances in R (R Core Team, 2021) to provide insight into diverse genotypes within some species. Thus, based on Jaccard and Bray-Curtis distances, we determined the optimal number of clusters based on the GAP statistic. Furthermore, we disentangled the number of genetic clusters in each species by Discriminant Analysis of Principal Components analysis (DAPC, Pritchard *et al*., 2000; Grünwald & Goss, 2011 and set the maximum number of possible cluster combinations to 10, and subsequently determined the optimal number based on the BIC statistic and the posterior membership probability (Grünwald & Goss, 2011).

### Illumina sequencing and whole-genome comparisons

The genomes of a subset of 22 Cuban strains that capture the diversity uncovered by the DArTseq analysis were sequenced using Illumina HiSeq PE150 (BGI, Hong Kong) and *de novo* assembled with Spades (Bankevich *et al*., 2012) version 3.13.0, with default settings. Contigs shorter than 500 bp were subsequently removed. To assess the assembly quality, the genome assemblies were analyzed with QUAST version 5.0.2 (Gurevich *et al*., 2013), and genome completeness was estimated based on the presence of conserved single-copy genes using BUSCO version 5.1.2 (Simão *et al*., 2015) with the sequencing odb10 database. To determine the relationship of the sequenced Cuban isolates with other banana-infecting *Fusarium* spp., we included whole-genome sequencing data of 58 previously sampled FWB pathogen isolates from different species and geographical locations (Table S4). To identify sequence variation, the sequencing reads were aligned against the chromosome-level genome assembly of the *F. odoratissimum* II5 (NRRL 54006) reference strain (van Westerhoven *et al*., 2023) using BWA-mem (v. 0.7.17) (Li, 2013). The SNPs of the 80 isolates compared with the reference genome assembly were called based on the GATK best practices (Auwera *et al*., 2013), using GATK4 (v. 4.2.0.0). To account for the presence/absence variation between the genomes, we excluded SNPs located on the highly variable genomic regions (chromosome 1, pos. 0-1.2 Mb, and chromosome 12) (van Westerhoven *et al*., 2023) as well as SNPs with any missing values. The maximum likelihood phylogenetics tree of the concatenated SNPs was inferred using RAxML (v. 8.2.12, - m GTRCAT). Nucleotide diversity values (π) of the samples were calculated with VCFtools (version 0.1.17) with haploid genome specification (Danecek *et al*., 2011).

## RESULTS

### Fusarium wilt of banana is widespread across Cuba

To obtain a nationwide overview of the incidence of FWB in Cuba, we completed a comprehensive survey from 2016 to 2018 that included all the provinces of the country and sampled bananas in large commercial fields, small farms, backyards, and roadsides. During the survey, plants with FWB symptoms were observed in 14 of the 15 provinces in Cuba, and in 43 of the 66 surveyed municipalities (Figure 1). In the province Las Tunas, we did not observe affected plants since most of the visited and investigated plantations were relatively young due to rejuvenation after the devastating hurricane ‘Irma’. In total, 130 isolates were obtained from discolored vascular pseudostem strands sampled from symptomatic banana plants (Table S2). The FWB-affected banana varieties comprised ‘Manzano vietnamita’ from the Pisang Awak subgroup (ABB), and the cooking banana varieties ‘Burro Criollo’, ‘Burro Enano’ (Dwarf), and ‘Burro CEMSA’ (Figure 1B) from the Bluggoe subgroup (ABB). Diseased plants showed the typical FWB symptoms but in ‘Burro CEMSA’ symptoms were occasionally limited to marginal leaf chlorosis and splitting of the pseudostem. In total, 37% and 43 % of the observed FWB cases were in fields of ‘Manzano vietnamita’ and ‘Burro CEMSA’, respectively, while the remaining 20% of FWB cases were in the other Bluggoe varieties (ABB). We did not observe any FWB in ‘Pisang Ceylan’ (AAB), ‘FHIA-18’ (AAAB), ‘FHIA-21’ (AAAB), and those of the Cavendish (AAA) and Plantain (AAB) subgroups, which are also widely cultivated in Cuba (Table S1), suggesting that these banana varieties are resistant to the *Fusarium* population on the island.

All isolates were initially identified as members of the FOSC based on morpho-cultural characteristics (Leslie & Summerell, 2006), like the production of oval to kidney-shaped microconidia in false heads on short monophialides, which were typically single-celled. Moreover, the different isolates produced chlamydospores which were single or in pairs and produced at hyphal ends or within the hyphae. Colonies on PDA plates varied in color, from white or pinkish to dark purple (Supporting Information Fig. S1). To further substantiate that the isolates belonged to the FOSC, all isolates were characterized by PCR following the protocols described by Edel *et al*. (2000), Dita *et al*. (2010), and Carvalhais *et al*. (2019), which confirmed that they belonged to FOSC and that they were either Race 1 or Race 2 strains. Importantly, all isolates tested negative with diagnostic TR4 primers (Supporting Information Table S3), which accords with the absence of FWB during the surveillance of the Cavendish and Plantain subgroups, which are resistant to Race 1 and Race 2.

### Fusarium isolates causing FWB in Cuba exclusively belong to Race 1 or Race 2

To corroborate the pathogenicity of the Cuban isolates to a subset of banana cultivars under standardized greenhouse conditions, we selected 11 isolates representative of various provinces and surveyed hosts, and tested them with differential cultivars for Race 1 (‘Gros Michel’), Race 2 (‘Burro CEMSA’), and Race 4 (‘Grand Naine’). All isolates tested caused characteristic FWB symptoms at 12 weeks after inoculation, on either Race 1 or Race 2 susceptible cultivars (Figure 2). The RDA scores in each banana cultivar varied according to the inoculated isolate. Race 1 isolates (cub0012, cub0035, cub0129, cub0130, and cub0140) caused RDA scores ranging from 27.9 to 61.3% in ‘Gros Michel’ but in ‘Burro CEMSA’ the RDA scores were lower than 16.1%. In contrast, ‘Gros Michel’ plants inoculated with Race 2 isolates (cub0022, cub0034, and cub0088) scored RDA values between 0 to 19.3%. However, these isolates caused RDA scores ranging from 25.2 to 64.4 % in ‘Burro CEMSA’ plants. Interestingly, three isolates (cub0043, cub0145, and cub0150) caused symptoms in both ‘Gros Michel’ and ‘Burro CEMSA’ plants, hence can neither be classified as Race 1 nor as Race 2 strains. Importantly, all inoculated ‘Grand Naine’ plants and the water controls remained free of FWB symptoms, which corroborated the PCR results, and confirms that Race 4 is likely not present in Cuba at the time of the survey.

**Fig. 2.**
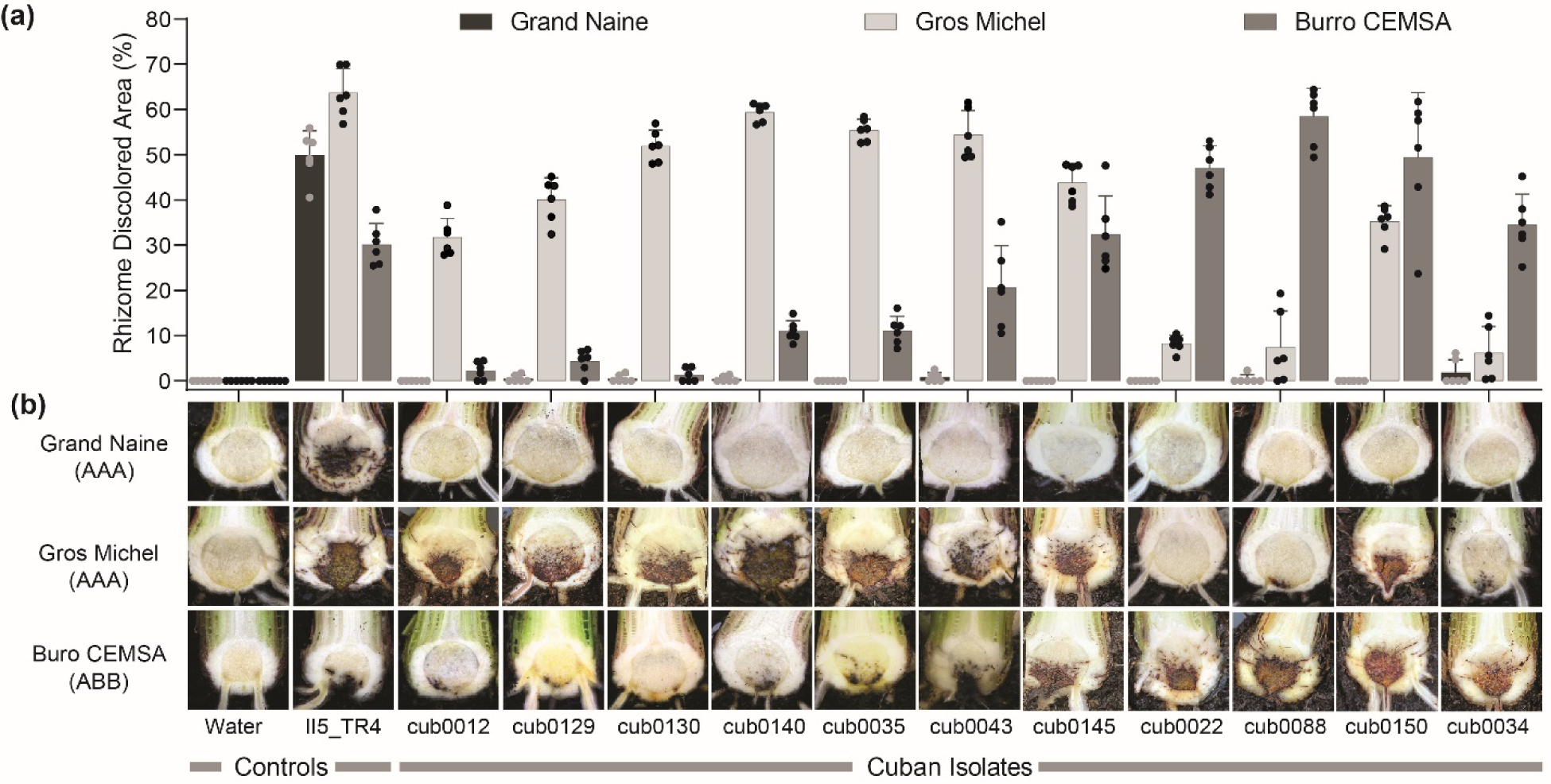
Cuban *Fusarium* strains are pathogenic in greenhouse infections assays and cause FWB on banana varieties that are susceptible to the so-called Race 1 (‘Gros Michel’) and Race 2 (‘Burro CEMSA’) strains. (A) Rhizome discolored area scores, calculated with ImageJ, were isolate-dependent and ranged from 0.2% to 61.3% in ‘Gros Michel’ and from 4.9% to 69.4% in ‘Burro CEMSA’. (B) Representative overview of FWB severity in the corms of banana cultivar ‘Grand Naine’, ‘Gros Michel’, and ‘Burro CEMSA’ 12 weeks after inoculation. Note that none of the Cuban strains caused FWB symptoms in the Cavendish cultivar ‘Grand Naine’.

### *The Cuban* Fusarium *populations that cause FWB contain both mating types*

The mating type frequency in fungal populations provides an indication of possible sexual recombination. Random mating usually results in a 1:1 ratio between the two mating type alleles in fungi with a bipolar, heterothallic mating system, and deviation from this ratio either indicates mating irregularities or even the absence of mating (Heitman *et al*., 2017). Although *Fusarium* spp. that cause FWB are thought to be asexual pathogens, they do carry the mating type idiomorphs *mat1-1* and *mat1-2* (Fourie *et al*., 2009) which have been used to study the evolution of various *formae speciales* of *F. oxysporum* (Fourie *et al*., 2009; Lievens *et al*., 2009). Thus, we sought to determine the *mat1-1* : *mat 1-2* ratio among the Cuban isolates. Similar to earlier findings in *Fusarium* spp. (Fourie *et al*., 2009; Visser *et al*., 2010; Ordóñez, 2018; Magdama *et al*., 2020), we observed that all isolates either contained the *mat1-1* or the *mat1-2* idiomorph. However, as expected from isolates that do not undergo regular sexual recombination, the ratio of *mat1-1* : *mat1-2* was 15:155 (Table S3). Thus, we conclude that even though both mating types occur across Cuba, and no strains with both mating type alleles were sampled, the skewed mating type frequency suggests the absence of a sexual cycle during the evolution of Cuban populations of the pathogen.

### Fusarium *spp. causing FWB in Cuba are genetically diverse*

Recently, the taxonomy of the *Fusarium* lineages that cause FWB was revised (Maryani *et al*., 2019). TR4 was considered as the new species *F. odoratissimum,* Race 2 strains belong to *F. phialophorum*, *F. tardicrescens* or *F. tardichlamydosporum,* while Race 1 strains are present in at least seven species (Maryani *et al*., 2019; Czislowski *et al*., 2021; Santos *et al*., 2022). To determine which species are present in Cuba and to uncover their genetic diversity, we generated genotyping-by-sequencing DArT markers across the 170 Cuban isolates and added these to DArT markers that were previously obtained from 243 additional isolates, comprising 210 isolates from the Americas and the 33 isolates from other regions worldwide (Supporting Information Table S4). In total, we obtained 33,298 high-quality DArT (presence/absence) markers with a mean call rate of 99.3% and 6,600 single-nucleotide polymorphisms (SNPs) across the 413 FOSC isolates, using *Fusarium fujikuroi* (CBS 221.76) as an outgroup.

We performed all analyses using either the DArT presence/absence markers or the SNPs dataset, and in both cases, the results were consistent (Fig. S2). Hierarchical clustering based on DArT presence/absence markers showed that the *Fusarium* isolates could be divided into FOSC clades 1, 2, and 3, and then into 10 independent lineages, which overall correspond with the phylogenetic species described by Maryani *et al*. (2019) (Fig. 3). In clade 1, isolates of *F. odoratissimum* were grouped with those of *F. purpurascens* and *F. phialophorum*. Isolates of *F. duoseptatum, F. grosmichelii, F. tardicrescens,* and *F. tardichlamydosporum* were grouped in clade 2. Clade 3 included a group of 29 *F. oxysporum* isolates from Brazil, Colombia, Nicaragua, and Perú (Table S3), which did not cluster with any of the representatives of the previously defined lineages. Finally, a small group of isolates formed clade 4, encompassing the *F. sangayamense* and *F. kalimantanense* lineages, which are endemic to Indonesia and non-pathogenic on ‘Gros Michel’ and ‘Grand Naine’ (Maryani *et al*., 2019), although the latter was recently reported to infect “Silk” (AAB) bananas in Brazil, which is indicative for Race 1 (Santos *et al*., 2022).

**Fig. 3.**
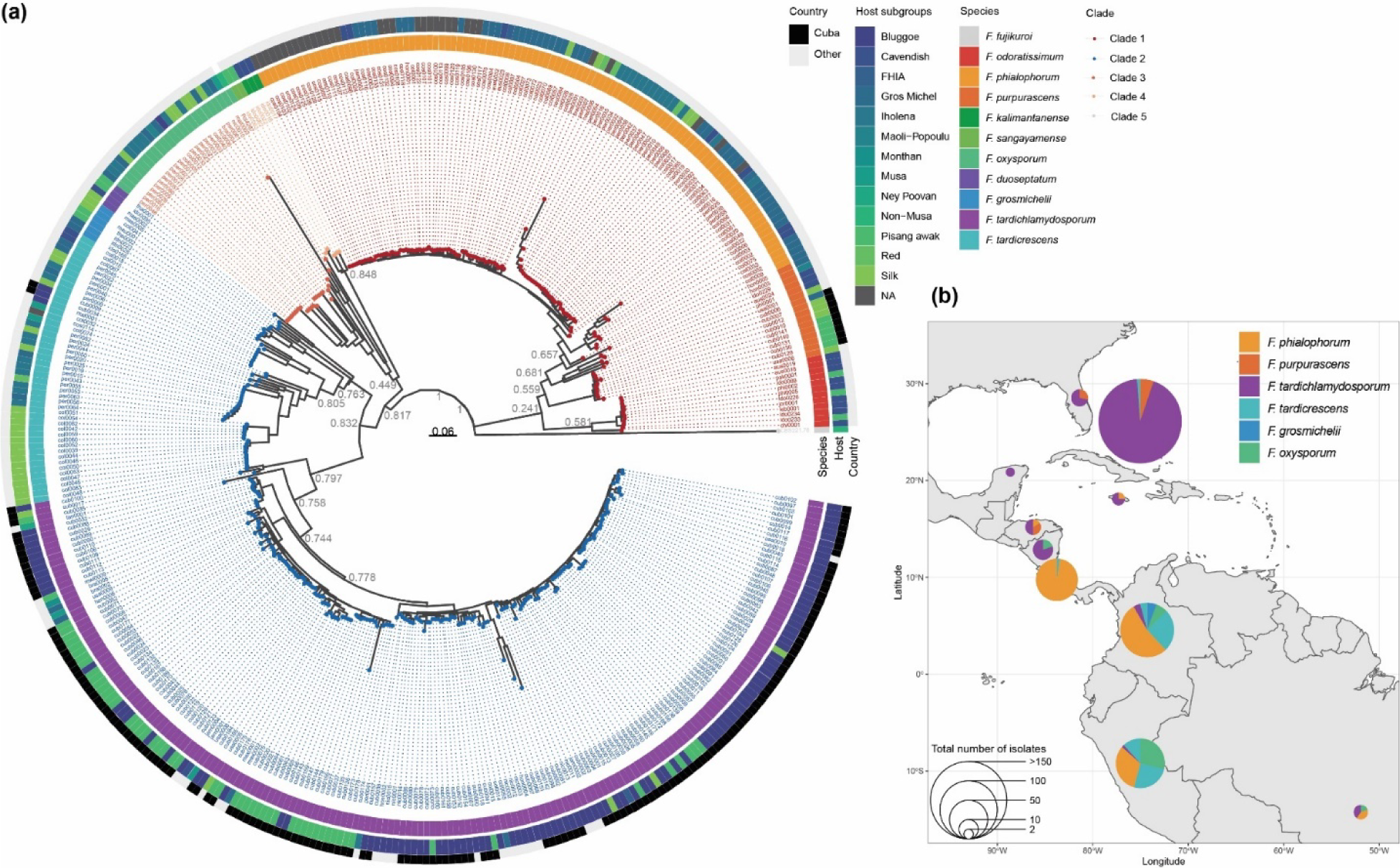
*Fusarium* isolates causing Fusarium wilt of banana (FWB) in the Americas are genetically diverse. A) Cuban isolates clustered in the lineages of *F. purpurascens*, *F. tardicrescens,* and *F. tardichlamydosporum* based on genotyping-by-sequencing (DArT) markers from a total of 414 isolates from 22 countries that were analyzed by the Neighbor-Joining method. The association of isolates to clades are shown as colored dots, red corresponds with clade 1, blue with clade 2, orange with clade 3, and yellow with clade 4. Species are represented as colored blocks in the inner circle, the middle circle corresponds with host and black or grey blocks in the outer circle indicate Cuban vs. non-Cuban isolates, respectively. The phylogenetic tree is rooted by *Fusarium fujikuroi* (CBS 221.76). B) The compositions of FWB-causing *Fusarium* spp. per country is variable. The species *F. tardichlamydosporum* and *F. purpurascens* are primarily present in the North Caribbean, whereas *F. phialophorum* and *F. tardicrescens* occur predominantly in Costa Rica and South American countries.

The great majority (95%) of the Cuban isolates grouped in the lineages corresponding with the species *F. tardichlamydosporum* and *F. tardicrescens* in clade 2, whereas only 5% grouped in the *F. purpurascens* lineage of clade 1. The *F. tardichlamydosporum* isolates include both Race 1 and Race 2 strains (Fig. 2, Table S3) and contain (94%) of the Cuban isolates, which affect ‘Burro Criollo’, ‘Burro CEMSA’, ‘Burro Enano’, ‘FHIA-03’, ‘Silk’ and ‘Pisang Awak’ (Table S2). Only two isolates belong to *F. tardicrescens* and were obtained from wilted ‘Burro Criollo’ plants in the Villa Clara province. The *F. purpurascens* isolates were phenotyped as Race 1 and affected ‘Gros Michel’, ‘Silk’, and ‘Pisang Awak’ bananas in the Central provinces Camagüey, Sancti Spiritus, and Villa Clara, as well as in the Western Mayabeque province.

Having determined the species composition of the Cuban FWB-causing *Fusarium* population, we sought to compare it with the diversity of the FWB pathogen populations in the Americas. We noticed that *F. tardichlamydosporum* and *F. purpurascens* are mostly present in the North Caribbean (from Nicaragua to Florida), whereas *F. phialophorum* and *F. tardicrescens* are primarily distributed across South America, but also in Costa Rica (Fig. 3). Furthermore, to calculate the genetic diversity within each species, we calculated the Jaccard and Bray-Curtis indices based on the DArT markers matrix and identified at least two subgroups in *F. tardichlamydosporum* (Figure S3). Both are present in Cuba, one mostly in the Northwest of the island and primarily associated with ‘Pisang Awak’, whereas the other is present in the Southeast and predominantly associated with Bluggoe cultivars (Fig. 4, Table S2), which suggests the presence of distinct genetic sub-clusters within the FWB causing species. To investigate the putative number of *Fusarium* genotypes present in the Americas, we performed a DAPC which revealed an optimum of six and a maximum of seven genetically distinct clusters (Fig. S3). Similarly, the number of genetic clusters exceeds the six species that we described for the American continent. As expected, based on the asexual nature of the FOSC, we did not observe any signs of admixture (Fig. 4), hence each isolate is assigned to a discrete genetic cluster. The species *F. grosmichelli, F. phialophorum, F. purpurascens,* and *F. oxysporum* form discrete clusters, but we also observed that Honduran isolates of *F. purpurascens* cluster closely with *F. phialophorum* isolates (Fig. 4), indicating a complex relationship between these species.

**Fig. 4.**
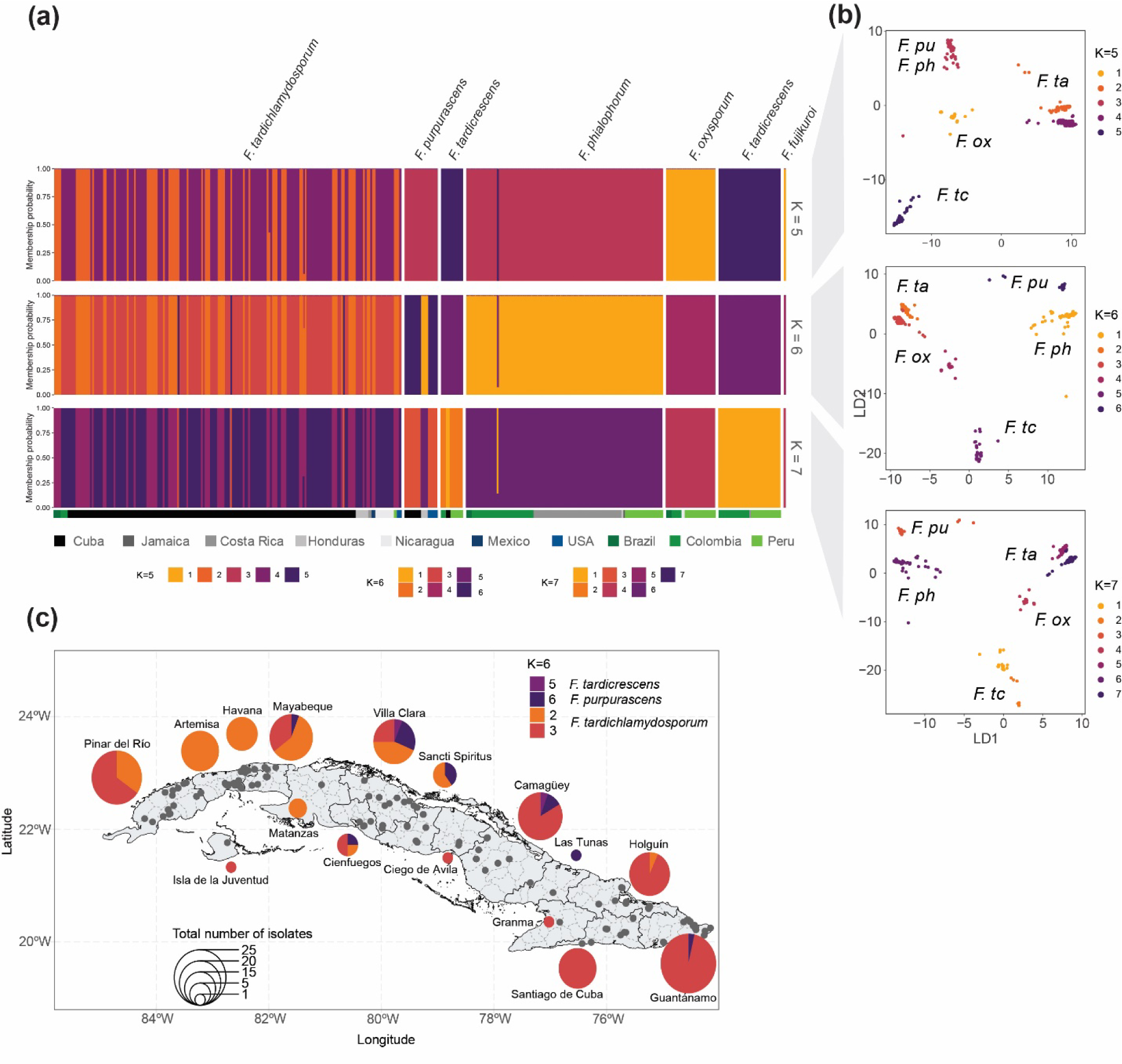
Discriminant analysis of principal components (DAPC) revealed the presence of an optimum of six genetically distinct clusters of *Fusarium* strains that cause Fusarium wilt of banana (FWB) in the Americas. a) Structure of FWB-causing *Fusarium* isolates according to DAPC for *K*=5-7, the bottom colors depict the country of origin. b) Clustering of the isolates based on *K*=5-7 highlights at least two genetic sub-clusters in *F*. *tardichlamydosporum.* Abbreviations: *F.ox = F. oxysporum* (isolates of Clade 3 that did not cluster with the representatives of the previously defined species)*, F.pu = F. purpurascens, F.ph = F. phialophorum, F.ta = F. tardichlamydosporum,* and *F.tc = F. tardicrescens*. c) Two genetically diverse sub-groups of *F*. *tardichlamydosporum* are present in Cuba. One group (light orange) is more abundant in the Center to the Northwest of the country, whereas the second group (maroon) is mostly present in the Southeast.

To further corroborate the DArT-based taxonomic classification, we compared whole-genome short-read sequences of 22 Cuban isolates together with 58 *Fusarium* isolates from a global panel of fully sequenced isolates (van Westerhoven *et al*., 2023; Table S5); the 22 Cuban isolates represent the range of diversity in terms of race, lineage, host, and geographic origin across the country. With this collection of isolates, we identified nearly 1,4 million high-quality SNPs and subsequently constructed a maximum-likelihood based phylogenetic tree. As to be expected, this analysis confirmed that FWB-causing isolates are grouped in 10 independent lineages, which were supported by high bootstrap values, and correspond to the assignments based on the DArT-based taxonomy. Moreover, we calculated the nucleotide diversity values (π) within each species and showed that π values per *Fusarium* lineage is generally low (Fig. 5), and consequently, our collective data strongly supports the phylogenetic species previously described by Maryani *et al*. (2019).

**Fig. 5.**
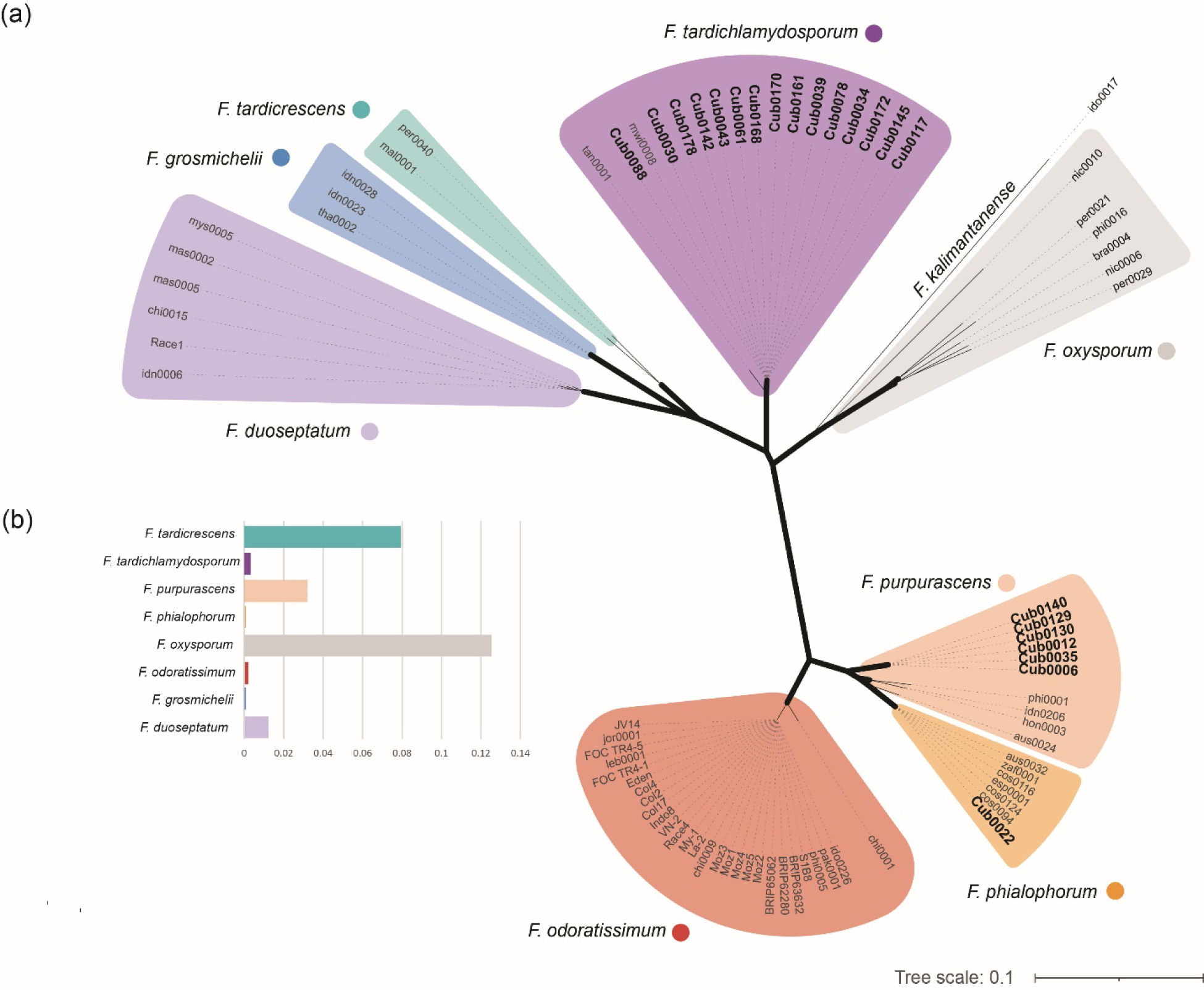
*Fusarium* isolates that cause Fusarium wilt of banana grouped in distinct lineages irrespective of their geographical origin. (a) The unrooted Maximum-Likelihood phylogenetic tree based on 1,391,348 high-quality SNPs, inferred from mapping sequence data of a subset of 80 isolates onto the reference genome of *F. odoratissimum* strain II5, shows that the identified groups are consistent with species naming mostly consistent with species naming. Thick branches indicate a bootstrap support of 100%. Cuban isolates clustered in the lineages of *F. purpurascens*, *F. phialophorum,* and *F. tardichlamydosporum*. Background shading delineates different *Fusarium* lineages. The isolate codes of those from Cuba are in bold. (b) Bar plot with the nucleotide diversity values (π) for each species determined between isolates for each *Fusarium* lineages.

## DISCUSSION

Roots, tubers, and bananas (including dessert bananas, plantains, and cooking bananas) are important food crops and valuable traded commodities in numerous developing countries where they play a critical role in ensuring food security and income (Scott, 2021). The importance of these crops for food security is due to their high yields and carbohydrate content which translates into a higher daily energy supply per cultivated hectare than cereals (Petsakos *et al*., 2019). Particularly, bananas are the most popular fruit worldwide and a primary staple food in tropical and subtropical regions where most bananas are produced (FAO, 2022). The importance of banana for food security is markedly relevant for East Africa, a region with the world’s highest per capita banana consumption (Akankwasa *et al*., 2021). Meanwhile for LAC, bananas are a very important cash crop with five countries of the region in the top 10 banana exporting nations (FAO, 2022). Furthermore, as in East Africa, bananas are also important as a staple food in local diets in LAC, with approximately 62% of the regional production consumed locally (Dita *et al*., 2013). In Cuba, bananas are produced in all provinces including the special municipality ‘Isla de la Juventud’ and are the most produced fruits nationally along with mango and guava, representing nearly 30% of all fruit and starchy root staples produced in the country (ONEI, 2022).

In the past, Cuban banana production was threatened by yellow Sigatoka and FWB, which destroyed the national export activities during the first half of the last century. However, when Cavendish bananas and plantains were adopted, FWB lost importance (Martínez-de la Parte *et al*., 2023). However, when Black Leaf Streak Disease, caused by *Pseudocercospora fijiensis* (Arango *et al*., 2016), entered the country in the 1990s, the overall assembly of the national banana acreage changed. Since then, areas grown with susceptible Cavendish and plantain cultivars were significantly reduced and replaced by more resistant FHIA hybrids and Bluggoe cultivars (Pérez-Vicente *et al*., 2002), and ‘Pisang Awak’ became popular due to its higher rusticity and semi-acid taste. These changes caused a reemergence of FWB in Cuba, which is corroborated by our analyses that demonstrated that this disease is widespread across the entire country.

We assembled a nationwide collection of 170 isolates from symptomatic bananas primarily from ‘Burro CEMSA’ and ‘Pisang Awak’, which are extensively grown in Cuba (Martínez-de la Parte *et al*., 2023). Although diagnostic PCRs and phenotyping assays showed that Race 1 and Race 2 prevailed in the collection, some isolates (isolates cub0043, cub0145, and cub0150) caused symptoms in both ‘Gros Michel’ and ‘Burro CEMSA’ which confirms previous field experiments in Cuba and greenhouse results with isolates from Brazil, East Africa, Florida, Nepal, and Puerto Rico (Ploetz & Churchill, 2011; Garcia *et al*., 2018; García-Bastidas, 2019; Martínez-de la Parte *et al*., 2023; Pant *et al*., 2023). Hence, the current race concept in FWB is inadequate because these isolates do not fit in the classical race nomenclature (i.e., Race 1, Race 2, and Race 4). We anticipate that more FWB races will be recognized with expanded phenotyping, including more banana accessions, similar to observations in *Fusarium* spp. causing Fusarium wilt in lettuce and watermelon (Zhou *et al*., 2009; Gilardi *et al*., 2017).

Phylogenetic analyses, based on several DNA-based techniques, have consistently shown that *Foc* isolates are divided into three major clades, which are further subdivided into eight to 10 lineages (Bentley *et al*., 1998; O’Donnell *et al*., 1998; Groenewald *et al*., 2006; Fourie *et al*., 2009; Mostert *et al*., 2017; Maryani *et al*., 2019). Furthermore, multi-locus genotyping and genome-by-sequencing using DArTseq showed that *Foc* is genetically complex comprising a suite of clearly defined lineages, including the various physiological races, that were consequently considered as new *Fusarium* species (Maryani *et al*., 2019). However, these analyses focused on Southeast Asia, which is the center of origin of *Musa*, but thus far there is no information about these species in the LAC context. Previous studies used DNA fingerprinting (Bentley *et al*., 1998), multigene phylogeny (Fourie *et al*., 2009; Maryani *et al*., 2019; Magdama *et al*., 2020), DArTseq (Mostert *et al*., 2022), or whole-genome sequence comparisons (Garcia-Bastidas *et al*., 2020; Acuña *et al*., 2021; Leiva *et al*., 2022; Reyes-Herrera *et al*., 2022), but included only a limited number of isolates from Latin America or only focused on individual countries (Magdama *et al*., 2020; Batista *et al*., 2022). Hence, no conclusions could yet be drawn on the phylogeography of FWB-causing fungi in LAC. Our study provides an unparalleled insight into the genetic diversity across a large set of 380 isolates obtained from 10 countries in LAC, including 170 isolates from Cuba. Therefore, contrary to the aforementioned studies, our data can resolve the structure of the FWB pathogens across this region. We conclude that FWB-causing fungi comprise five to seven genetic clusters, as isolates of *F. tardichlamydosporum* can be split into two distinct genetic sub-groups suggesting further population differentiation according to the type of banana cultivar from which they were sampled. Furthermore, some *F. purpurascens* isolates cluster closely with *F. phialophorum* in the phylogenetic tree based on DArT presence/absence markers, despite their position in the PCA and the phylogenetic tree based on DArT SNPs. Hence, the phylogenetic relationship between both species is intricate and requires further analysis. Although a recent report proposed that sexual reproduction occurs in the related chickpea pathogen *F. oxysporum* f. sp. *ciceris* (Fayyaz *et al*. 2023) our data on the ratio between the *mat 1-1* and *mat 1-2* idiomorphs neither support such a hypothesis in FWB-causing *Fusarium* spp. nor explain the aforementioned observations or indicate a sex-driven speciation. On the contrary, the mating type allele ratios are in accordance with previous reports (Fourie *et al*., 2009; Visser *et al*., 2010; Ordoñez, 2018; Magdama *et al*., 2020), suggesting the absence of sexual reproduction between *Fusarium* isolates causing FWB. Taken together, our analyses provide a high-resolution and genome-wide dataset to describe genetic diversity in FWB*-*causing *Fusarium* spp. in an unparalleled manner that matches recent genome-wide analyses (van Westerhoven *et al*., 2023: Maryani *et al*., 2023).

The genotyping data showed that the Cuban isolates are grouped in Clades 1 and 2 and in three lineages of which none clustered with the *F. odoratissimum* lineage, which is in line with the pathogenicity tests and molecular diagnostics; no strain caused disease in Cavendish and the PCR data that were all negative for TR4 (Dita *et al*., 2010). Nevertheless, Cuban banana production is threatened by the recent expansion of TR4 in Latin America (Garcia-Bastidas *et al*., 2020; Acuña *et al*., 2021; Herrera *et al*., 2023; Martínez de la Parte *et al*., 2023). Overall, phylogenies based on DArT and whole-genome SNPs are in accord for 99.8% of the isolates, and thus we conclude that FWB pathogens in Cuba belong to the species *F. purpurascens, F. tardichlamydosporum,* and *F. phialophorum*. The DAPC analysis showed that Cuban isolates of a single species are closely related with isolates of the same species from different geographies, compared to their relationship with other Cuban isolates of other *Fusarium* spp., which suggests several independent introductions of *Fusarium* since the inception of banana cultivation. Bananas arrived in Cuba in 1529 from Hispaniola Island, now Dominican Republic, and soon became one of the most important staple foods for the population (García, 2001). Over time, several banana cultivars from different origins were independently introduced into Cuba, such as Gros Michel from Martinique, Grand Naine from Panama, and plantains from the Dominican Republic and Saint Lucia (Marin *et al*., 1998; García, 2008; Alvarez, 2011). Additionally, movement of banana planting material in LAC occurred (Marin *et al*., 1998), which may explain the observed close relationship and similar species composition of *Fusarium* populations in Central America and the Caribbean compared to populations from South America (Brazil, Colombia, and Perú). It is also possible that the introduction of planting material from different producing areas facilitated the introduction of genetically diverse pathogen strains. This is in line with the currently observed diversity in Cuba, which is also illustrated by the similarity with isolates from Cuba and Florida, USA. Both countries harbor a unique *Fusarium* population comprising isolates of VCG01210 that is only found in these countries and in the Cayman Islands (Ploetz, 1990; Pérez-Vicente, 2004; Mostert *et al*., 2017), which could be explained by the historical ties between Cuba and the USA through trade and migration. FWB was first described in Cuba in 1910 and occurred in Florida decades later (1986), suggesting that similar *Fusarium* genotypes – characterized as VCG01210 - may have been introduced from Cuba via infected Silk banana plants. Therefore, our analyses also underscore human mobility and behavior as critical factors in previous and recent outbreaks of FWB (Garcia-Bastidas *et al*., 2020; Acuña *et al*., 2021; Herrera *et al*., 2023), which threaten banana production across LAC.

## ACKNOWLEDGEMENTS

EMP was supported by a NUFFIC PhD scholarship, grant number EPS 2016-02. GHJK & HJGM were supported by the Dutch Dioraphte Foundation. DET acknowledges his PhD fellowship from the Consejo Nacional de Ciencia y Tecnología from México. This work was supported in part by grants from the research projects P131LH003061 and P131LH003060 of the Cuban Program of Animal and Plant Health of the Cuban Ministry of Agriculture. The funders had no role in study design, data collection and analysis, decision to publish, or preparation of the manuscript.

## AUTHOR CONTRIBUTION

GHJK and LPV conceived and supervised the project. EMP collected data and conducted the analyses. LPV, DET, and ACW contributed to the collected data. DET, ACW, and MFS contributed to the analyses. EMP, DET, and GHJK wrote the manuscript with contributions from MFS. All authors (EMP, LPV, DET, ACW, HJGM, MFS, and GHJK) edited and approved the manuscript.

