## Supplementary material for "A deep genetic analysis of banana Fusarium wilt pathogens of Cuba in a Latin American and Caribbean diversity landscape": Fig. S1

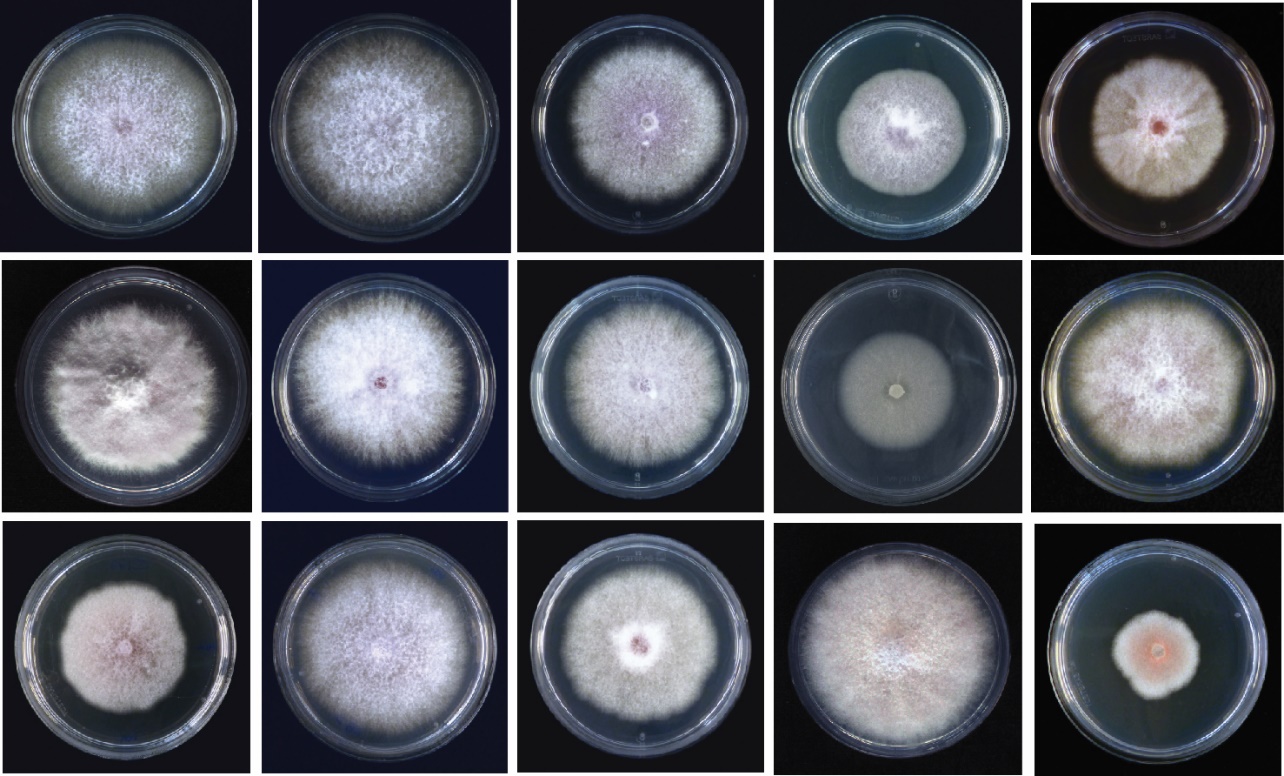

**Fig. S1** Cultural characteristics of Cuban isolates of different *Fusarium* species that cause FWB. Colonies of different isolates varied in color, texture and colony diameter after one week incubation in PDA plates at 25°C.
