## Supplementary material for "A deep genetic analysis of banana Fusarium wilt pathogens of Cuba in a Latin American and Caribbean diversity landscape": Fig. S2

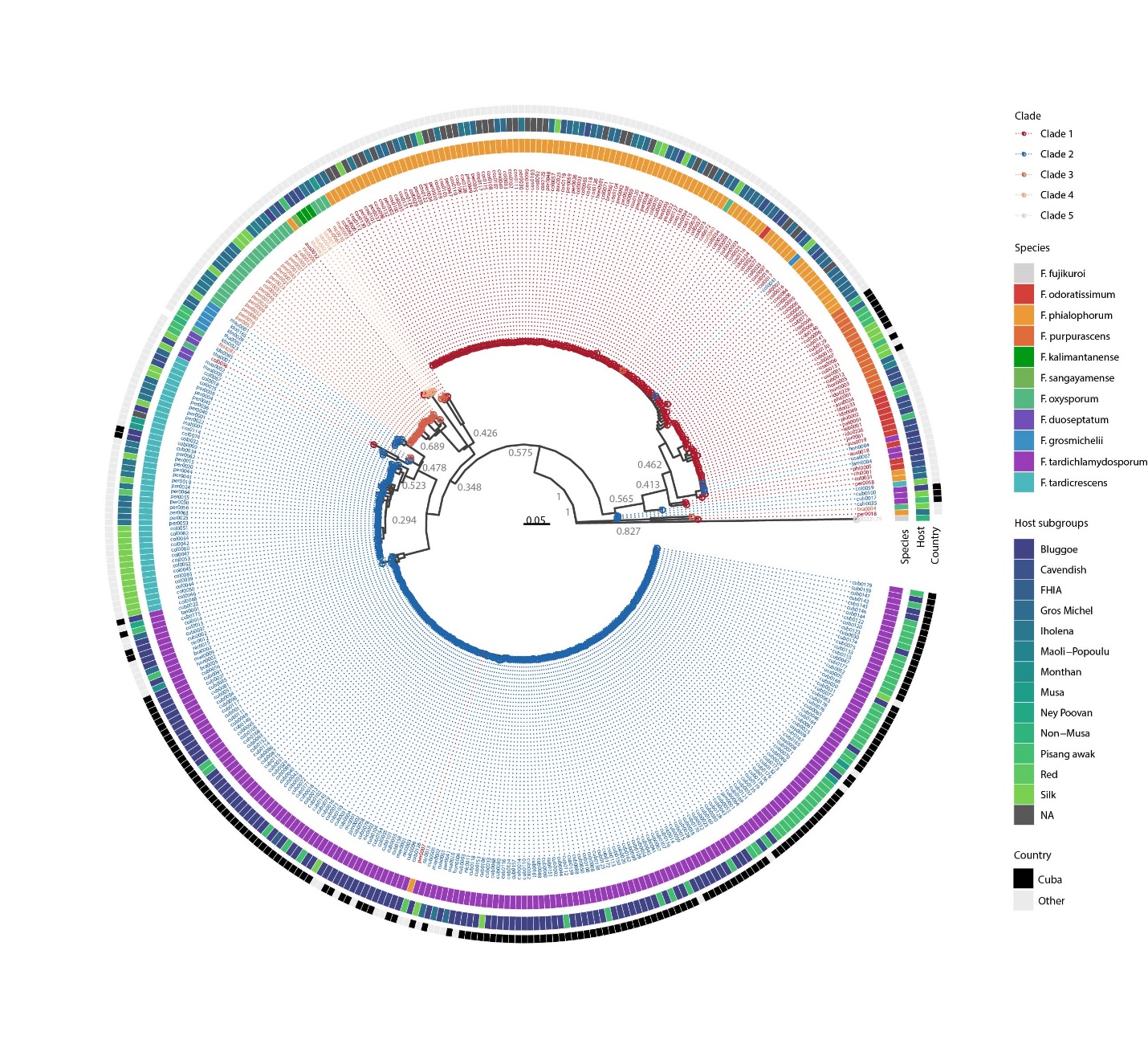

**Fig. S2** *Fusarium* isolates causing Fusarium wilt of banana (FWB) in the Americas are genetically diverse as shown by a Neighbor-Joining tree inferred from single-nucleotide polymorphisms derived from the DArT markers. The association of isolates to clades are shown as colored dots, red corresponds with clade 1, blue with clade 2, orange with clade 3, and yellow with clade 4. *Fusarium* species as defined by Maryani *et al*. (2019) are shown as colored blocks in the inner circle, the middle circle corresponds with host, and black or grey blocks in the outer circle indicate Cuban vs. non-Cuban isolates, respectively. The entire tree is rooted by *Fusarium fujikuroi* (CBS 221.76).
