## Supplementary material for "A deep genetic analysis of banana Fusarium wilt pathogens of Cuba in a Latin American and Caribbean diversity landscape": Fig. S3

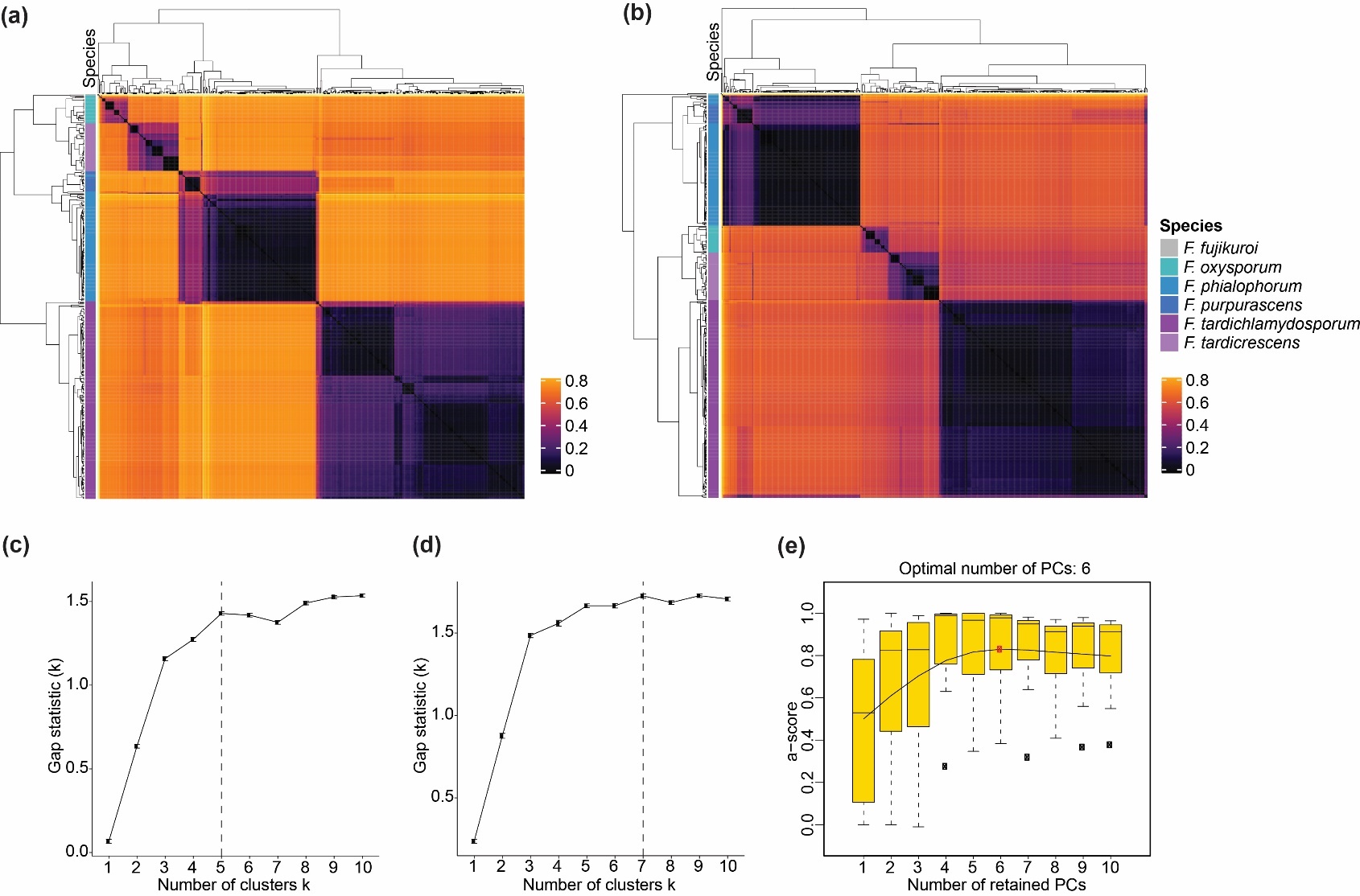

**Fig. S3** American strains of the causal agent of FWB are grouped into five to seven genetically distinct groups. A-B) Heatmap of the Jaccard distance (A) and Bray-Curtis dissimilarity (B) between 380 *Fusarium* strains. C-E) The optimal number of clusters was determined according to Jaccard distance (C), Bray-Curtis distance (D), or a Discriminant Analysis of Principal Components (E), which aims to reduce variation within groups and to maximize variation between groups.
