## Supplementary material for "A deep genetic analysis of banana Fusarium wilt pathogens of Cuba in a Latin American and Caribbean diversity landscape": Table S1

**Table S1**. Composition of the total banana acreage (180596 ha.) of Cuba according to the last national survey (Plant Health Directorate of the Cuban Ministry of Agriculture, DSV 2014, not published).

| **TYPE** | **Subgroup** | **Variety** | **Genome**  **composition** | | **Planted Area (ha)** |
| --- | --- | --- | --- | --- | --- |
| Dessert | Cavendish | Gran Naine | AAA | 6847 | |
| Dessert | Cavendish | Robusta (Valery) | AAA | 12223 | |
| Dessert | Gros Michel | Gros Michel | AAA | 513 | |
| Dessert | Cavendish | Bungulan | AAA | 85 | |
| Dessert | Cavendish | Giant Cavendish | AAA | 65 | |
| Dessert | Cavendish | Dwarf Cavendish | AAA | 42 | |
| Dessert | Ibota | Yangambi Km5 | AAA | 3 | |
| Dessert | FHIA hybrids | FHIA-02 | AAAA | 3 | |
| Dessert | FHIA hybrids | SH-3436-9 | AAAA | 1 | |
| Dessert | FHIA hybrids | FHIA-23 | AAAA | 48 | |
| Dessert | FHIA hybrids | FHIA-01 | AAAB | 274 | |
| Dessert | FHIA hybrids | FHIA-18 | AAAB | 6811 | |
| Dessert | Mysore | Pisang ceylan | AAB | 409 | |
| Dessert | Silk | Manzano INIVIT | AAB | 327 | |
| Dessert | Silk | Manzano | AAB | 6 | |
| Dessert | Pisang Awak | Manzano vietnamita | ABB | 1043 | |
| Cooking | Bluggoe | Burro Cemsa | ABB | 77801 | |
| Cooking | Bluggoe | Burro Criollo | ABB | 858 | |
| Cooking | Bluggoe | Burro Enano | ABB | 415 | |
| Cooking | FHIA hybrids | FHIA-25 | AAB | 371 | |
| Cooking | Bluggoe | Other Bluggoe | ABB | 39 | |
| Cooking | FHIA hybrids | FHIA-03 | AABB | 2 | |
| Plantain | Plantain | INIVIT PV-0630 | AAB | 1428 | |
| Plantain | Plantain | CEMSA3/4 | AAB | 37087 | |
| Plantain | Plantain | Macho3/4 | AAB | 19054 | |
| Plantain | Plantain | Enano Guantanamero | AAB | 6236 | |
| Plantain | FHIA hybrids | FHIA-21 | AAAB | 8549 | |
| Plantain | FHIA hybrids | FHIA-20 | AAAB | 2 | |
| Plantain | Plantain | Other Plantain | AAB | 50 | |
| Other | Others | Others | - | 5 | |
