## Supplementary material for "A deep genetic analysis of banana Fusarium wilt pathogens of Cuba in a Latin American and Caribbean diversity landscape": Table S2

**Table S2**. Origen of the Cuban *Fusarium* isolates, that cause Fusarium wilt of banana, characterized in this study.

| **DArT id** | **Isolate Code^1^** |  | **Geographical origin** | | | |  | **Host** | | | **Survey** |
| --- | --- | --- | --- | --- | --- | --- | --- | --- | --- | --- | --- |
|  |  |  | **Province** | **Municipality** | **coordinates** | **MASL^2^** |  | **Cultivar** | **Subgroup** | **genotype** |  |
| cub0001 | IJ-1 |  | Isla de la Juventud | Isla de la Juventud | 21.761454, -82.736489 | 39 |  | Burro CEMSA | Bluggoe | ABB | 1996-2000 |
| cub0002 | SC-3 |  | Santiago de Cuba | Palma Soriano | 20.226192, -75.989683 | 133 |  | Burro criollo | Bluggoe | ABB | 1996-2000 |
| cub0003 | C.Esm1.3 |  | Camaguey | Esmeralda | n.d. | n.d. |  | Burro criollo | Bluggoe | ABB | 1996-2000 |
| cub0004 | Cca1.1 |  | Camaguey | Camaguey | n.d. | n.d. |  | Manzano | Silk | AAB | 1996-2000 |
| cub0005 | R.Go |  | Havana | Playa | n.d. | n.d. |  | Manzano | Silk | AAB | 1996-2000 |
| cub0006 | Cub9 |  | Villa Clara | n.d. | n.d. | n.d. |  | Gros Michel | Gros Michel | AAA | - |
| cub0007 | M3 |  | Villa Clara | n.d. | n.d. | n.d. |  | Manzano | Silk | AAB | - |
| cub0008 | C.Esm1.1 |  | Camaguey | Esmeralda | 21.851445, -78.116708 | 37 |  | Burro criollo | Bluggoe | ABB | 1996-2000 |
| cub0009 | CGua 1.1 |  | Camaguey | Guaimaro | 21.049077, -77.353706 | 83 |  | Burro criollo | Bluggoe | ABB | 1996-2000 |
| cub0010 | C.V 2.1 |  | Camaguey | Vertientes | 21.270623, -78.155012 | 58 |  | Manzano | Silk | AAB | 1996-2000 |
| cub0011 | ESB-1 |  | Camaguey | Esmeralda | 21.843757, -78.123587 | 38 |  | Burro criollo | Bluggoe | ABB | 1996-2000 |
| cub0012 | Cam3 |  | Camaguey | Camaguey | 21.397476, -77.929907 | 95 |  | Manzano | Silk | AAB | 1996-2000 |
| cub0013 | SC-2 |  | Santiago de Cuba | Palma Soriano | 20.230811, -76.011663 | 136 |  | FHIA-03 | FHIA hybrid | AABB | 1996-2000 |
| cub0014 | GuCUM |  | Guantanamo | Maisi | 20.191915, -74.251045 | 380 |  | Burro CEMSA | Bluggoe | ABB | 1996-2000 |
| cub0015 | GuBaCu |  | Guantanamo | Maisi | 20.214089, -74.410481 | 72 |  | Burro Criollo | Bluggoe | ABB | 1996-2000 |
| cub0016 | GuMa |  | Guantanamo | Maisi | 20.244781, -74.153092 | 17 |  | Burro Criollo | Bluggoe | ABB | 1996-2000 |
| cub0017 | TUPP-1 |  | Las Tunas | Puerto Padre | 21.183148, -76.621987 | 18 |  | Manzano | Silk | AAB | 1996-2000 |
| cub0018 | GuBaPo |  | Guantanamo | Baracoa | 20.291000, -74.464333 | 29 |  | Burro Criollo | Bluggoe | ABB | 1996-2000 |
| cub0020 | MoGu 1 |  | Artemisa | Guira de Melena | n.d. | n.d. |  | Manzano vietnamita | Pisang Awak | ABB | 1996-2000 |
| cub0021 | MoGu 3 |  | Artemisa | Guira de Melena | n.d. | n.d. |  | Manzano vietnamita | Pisang Awak | ABB | 1996-2000 |
| cub0022 | WaHa |  | La Habana | Boyeros | 23.008254, -82.419289 | 71 |  | Burro CEMSA | Bluggoe | ABB | 1996-2000 |
| cub0023 | PaGu |  | Artemisa | Alquízar | 22.816530, -82.622965 | 23 |  | Manzano vietnamita | Pisang Awak | ABB | 1996-2000 |
| cub0024 | AlqHa 1 |  | Artemisa | Alquízar | n.d. | NA |  | Manzano vietnamita | Pisang Awak | ABB | 1996-2000 |
| cub0025 | GuHa |  | Artemisa | Guira de Melena | 22.780703, -82.529753 | 12 |  | Manzano vietnamita | Pisang Awak | ABB | 1996-2000 |
| cub0026 | PeBa |  | Guantanamo | Baracoa | 20.352072, -74.630209 | 46 |  | Burro CEMSA | Bluggoe | ABB | 1996-2000 |

**Table S2**. (Continue)

| **DArT id** | **Isolate Code^1^** |  | **Geographical origin** | | | |  | **Host** | | | **Survey** |
| --- | --- | --- | --- | --- | --- | --- | --- | --- | --- | --- | --- |
|  |  |  | **Province** | **Municipality** | **coordinates** | **MASL^2^** |  | **Cultivar** | **Subgroup** | **genotype** |  |
| cub0027 | ToaGu |  | Guantanamo | Baracoa | n.d. | NA |  | Burro criollo | Bluggoe | ABB | 1996-2000 |
| cub0028 | Bar 1 |  | Guantanamo | Baracoa | 20.3606683, -74.5144867 | 29 |  | Burro criollo | Bluggoe | ABB | 1996-2000 |
| cub0029 | MaHo 1 |  | Holguin | Mayarí | n.d. | n.d. |  | Burro CEMSA | Bluggoe | ABB | 1996-2000 |
| cub0030 | BaHo 1 |  | Holguin | Banes | 20.958913, -75.720306 | 36 |  | Burro criollo | Bluggoe | ABB | 1996-2000 |
| cub0031 | BaHo 2 |  | Holguin | Banes | 20.964810, -75.734556 | 41 |  | Burro criollo | Bluggoe | ABB | 1996-2000 |
| cub0032 | SaHo |  | Holguin | Sagua de Tánamo | 20.625285, -75.230387 | 7 |  | Burro CEMSA | Bluggoe | ABB | 1996-2000 |
| cub0033 | Bar 3 |  | Guantanamo | Baracoa | 20.356750, -74.516493 | 19 |  | Manzano | Silk | AAB | 1996-2000 |
| cub0034 | R-2 |  | Villa Clara | Santo Domingo | 22.586179, -80.219636 | 48 |  | Burro criollo | Bluggoe | ABB | 1996-2000 |
| cub0035 | JCien |  | Cienfuegos | Cienfuegos | 22.1332, -80.3882 | 33 |  | Manzano vietnamita | Pisang Awak | ABB | 2016-2018 |
| cub0037 | CaVC |  | Villa Clara | Caibarién | 22.4192, -79.4228 | 18 |  | Manzano vietnamita | Pisang Awak | ABB | 2016-2018 |
| cub0038 | ToBaGu |  | Guantanamo | Baracoa | 20.386083, -74.569061 | 22 |  | Burro CEMSA | Bluggoe | ABB | 2016-2018 |
| cub0039 | NarBaGu |  | Guantanamo | Baracoa | 20.3906, -74.5489 | 10 |  | Burro CEMSA | Bluggoe | ABB | 2016-2018 |
| cub0040 | PGGua1 |  | Santiago de Cuba | Guamá | 19.991499, -76.272205 | 11 |  | Burro Criollo | Bluggoe | ABB | 2016-2018 |
| cub0041 | PGGua2 |  | Santiago de Cuba | Guamá | 19.991499, -76.272205 | 11 |  | Manzano vietnamita | Pisang Awak | ABB | 2016-2018 |
| cub0042 | BaGra |  | Granma | Bayamo | 20.3491, -76.8307 | 40 |  | Burro CEMSA | Bluggoe | ABB | 2016-2018 |
| cub0043 | PunBra1 |  | La Habana | La Lisa | 23.02041, -82.489242 | 60 |  | Manzano vietnamita | Pisang Awak | ABB | 2016-2018 |
| cub0044 | PunBra2 |  | La Habana | La Lisa | 23.019867, -82.487798 | 61 |  | Manzano vietnamita | Pisang Awak | ABB | 2016-2018 |
| cub0045 | BatMay1 |  | Mayabeque | Batabanó | 22.7801, -82.2996 | 33 |  | Burro enano | Bluggoe | ABB | 2016-2018 |
| cub0046 | BatMay2 |  | Mayabeque | Batabanó | 22.7806, -82.2984 | 33 |  | Manzano vietnamita | Pisang Awak | ABB | 2016-2018 |
| cub0047 | Min1Art |  | Artemisa | Artemisa | 22.77132, -82.723665 | 14 |  | Manzano vietnamita | Pisang Awak | ABB | 2016-2018 |
| cub0048 | MaySC1 |  | Santiago de Cuba | Segundo Frente | 20.424554, -75.525176 | 190 |  | Burro Criollo | Bluggoe | ABB | 2016-2018 |
| cub0049 | MaySC2 |  | Santiago de Cuba | Segundo Frente | 20.414686, -75.527427 | 177 |  | Burro Criollo | Bluggoe | ABB | 2016-2018 |
| cub0050 | CumCF1 |  | Cienfuegos | Cumanayagua | 22.054055, -80.320488 | 11 |  | Burro CEMSA | Bluggoe | ABB | 2016-2018 |
| cub0051 | CumCF2 |  | Cienfuegos | Cumanayagua | 22.064173, -80.303424 | 27 |  | Burro CEMSA | Bluggoe | ABB | 2016-2018 |
| cub0052 | BoyHa |  | La Habana | Boyeros | 23.043357, -82.366863 | 82 |  | Manzano vietnamita | Pisang Awak | ABB | 2016-2018 |

**Table S2**. (Continue)

| **DArT id** | **Isolate Code^1^** |  | **Geographical origin** | | | |  | **Host** | | | **Survey** |
| --- | --- | --- | --- | --- | --- | --- | --- | --- | --- | --- | --- |
|  |  |  | **Province** | **Municipality** | **coordinates** | **MASL^2^** |  | **Cultivar** | **Subgroup** | **genotype** |  |
| cub0053 | CotHa |  | La Habana | Cotorro | 23.065313, -82.255912 | 86 |  | Manzano vietnamita | Pisang Awak | ABB | 2016-2018 |
| cub0054 | GuaHa1 |  | La Habana | Guanabacoa | 23.066299, -82.253049 | 100 |  | Manzano vietnamita | Pisang Awak | ABB | 1996-2000 |
| cub0055 | C.Esm1.2 |  | Camaguey | Esmeralda | 21.851445, -78.116708 | 37 |  | Burro criollo | Bluggoe | ABB | 1996-2000 |
| cub0056 | C.Esm2.1 |  | Camaguey | Esmeralda | 21.851445, -78.116708 | 37 |  | Burro criollo | Bluggoe | ABB | 1996-2000 |
| cub0057 | ESB-5 |  | Camaguey | Esmeralda | 21.843757, -78.123587 | 38 |  | Burro criollo | Bluggoe | ABB | 1996-2000 |
| cub0058 | SCGu |  | Santiago de Cuba | Guamá | 19.968256, -76.433774 | 11 |  | Burro criollo | Bluggoe | ABB | 1996-2000 |
| cub0059 | Bar 2 |  | Guantanamo | Baracoa | 20.3606683, -74.5144867 | 29 |  | Burro criollo | Bluggoe | ABB | 1996-2000 |
| cub0060 | ColCA |  | Ciego de Avila | Baragua | 21.77283, -78.61835 | 18 |  | Burro enano | Bluggoe | ABB | 2016-2018 |
| cub0061 | Min2Art |  | Artemisa | Artemisa | 22.772401, -82.723595 | 14 |  | Manzano vietnamita | Pisang Awak | ABB | 2016-2018 |
| cub0063 | Art1Art-1 |  | Artemisa | Artemisa | 22.790584, -82.720201 | 29 |  | Manzano vietnamita | Pisang Awak | ABB | 2016-2018 |
| cub0064 | Art1Art-2 |  | Artemisa | Artemisa | 22.790584, -82.720201 | 29 |  | Manzano vietnamita | Pisang Awak | ABB | 2016-2018 |
| cub0065 | Art2Art-1 |  | Artemisa | Artemisa | 22.790572, -82.719580 | 29 |  | Manzano vietnamita | Pisang Awak | ABB | 2016-2018 |
| cub0067 | MelArt1.1 |  | Mayabeque | Melena del Sur | 22.798744, -82.196981 | 43 |  | Burro Cemsa | Bluggoe | ABB | 2016-2018 |
| cub0068 | MelArt1.2 |  | Mayabeque | Melena del Sur | 22.798744, -82.196981 | 43 |  | Burro Cemsa | Bluggoe | ABB | 2016-2018 |
| cub0069 | Front1Art1 |  | Artemisa | Artemisa | 22.778835, -82.721711 | 33 |  | Manzano vietnamita | Pisang Awak | ABB | 2016-2018 |
| cub0070 | Front1Art2 |  | Artemisa | Artemisa | 22.778835, -82.721711 | 33 |  | Manzano vietnamita | Pisang Awak | ABB | 2016-2018 |
| cub0071 | Front2Art |  | Artemisa | Artemisa | 22.779190, -82.721217 | 33 |  | Manzano vietnamita | Pisang Awak | ABB | 2016-2018 |
| cub0072 | FloCam1 |  | Camaguey | Florida | 21.472639, -78.315359 | 44 |  | Burro enano | Bluggoe | ABB | 2016-2018 |
| cub0073 | CesCam1 |  | Camaguey | Cespedes | 21.633698, -78.272799 | 42 |  | Manzano vietnamita | Pisang Awak | ABB | 2016-2018 |
| cub0075 | LisHa2 |  | La Habana | La Lisa | 23.039357, -82.475418 | 43 |  | Manzano vietnamita | Pisang Awak | ABB | 2016-2018 |
| cub0076 | TolHa |  | La Habana | Marianao | 23.054845, -82.420970 | 52 |  | Manzano vietnamita | Pisang Awak | ABB | 2016-2018 |
| cub0077 | AldHa |  | La Habana | Boyeros | 23.053978, -82.38377 | 68 |  | Manzano vietnamita | Pisang Awak | ABB | 2016-2018 |
| cub0078 | FloCam2 |  | Camaguey | Florida | 21.47199, -78.313663 | 51 |  | Burro criollo | Bluggoe | ABB | 2016-2018 |
| cub0079 | FloCam3 |  | Camaguey | Florida | 21.47199, -78.313363 | 51 |  | Burro criollo | Bluggoe | ABB | 2016-2018 |
| cub0080 | FloCam4 |  | Camaguey | Florida | 21.472675, -78.313992 | 51 |  | Burro criollo | Bluggoe | ABB | 2016-2018 |

**Table S2**. (Continue)

| **DArT id** | **Isolate Code^1^** |  | **Geographical origin** | | | |  | **Host** | | | **Survey** |
| --- | --- | --- | --- | --- | --- | --- | --- | --- | --- | --- | --- |
|  |  |  | **Province** | **Municipality** | **coordinates** | **MASL^2^** |  | **Cultivar** | **Subgroup** | **genotype** |  |
| cub0081 | CCam1 |  | Camaguey | Camaguey | 21.471365, -77.726914 | 83 |  | Burro enano | Bluggoe | ABB | 2016-2018 |
| cub0082 | CCam2 |  | Camaguey | Camaguey | 21.471778, -77.726673 | 83 |  | Burro enano | Bluggoe | ABB | 2016-2018 |
| cub0084 | CCam3.2 |  | Camaguey | Camaguey | 21.471907, -77.726643 | 83 |  | Burro enano | Bluggoe | ABB | 2016-2018 |
| cub0085 | MaHo2 |  | Holguin | Mayarí | 20.683855, -75.654668 | 12 |  | Burro Cemsa | Bluggoe | ABB | 2016-2018 |
| cub0086 | MaHo3 |  | Holguin | Mayarí | 20.683829, -75.654668 | 12 |  | Burro Cemsa | Bluggoe | ABB | 2016-2018 |
| cub0087 | MaHo4 |  | Holguin | Mayarí | 20.683874, -75.654712 | 12 |  | Burro Cemsa | Bluggoe | ABB | 2016-2018 |
| cub0088 | MaHo5 |  | Holguin | Mayarí | 20.709818, -75.647436 | 9 |  | Burro Cemsa | Bluggoe | ABB | 2016-2018 |
| cub0089 | MaHo6 |  | Holguin | Mayarí | 20.710014, -75.647722 | 9 |  | Burro Cemsa | Bluggoe | ABB | 2016-2018 |
| cub0090 | MaHo7 |  | Holguin | Mayarí | 20.709832, -75.6478 | 9 |  | Burro Cemsa | Bluggoe | ABB | 2016-2018 |
| cub0091 | PaHo1 |  | Holguin | Sagua de Tánamo | 20.615969, -75.248517 | 12 |  | Burro Cemsa | Bluggoe | ABB | 2016-2018 |
| cub0092 | PaHo2 |  | Holguin | Sagua de Tánamo | 20.615971, -75.248511 | 12 |  | Burro Cemsa | Bluggoe | ABB | 2016-2018 |
| cub0093 | SaHo2 |  | Holguin | Sagua de Tánamo | 20.589202, -75.252072 | 16 |  | Burro Criollo | Bluggoe | ABB | 2016-2018 |
| cub0095 | SaHo3.2 |  | Holguin | Frank Pais | 20.62353, -75.232295 | 10 |  | Burro Cemsa | Bluggoe | ABB | 2016-2018 |
| cub0096 | SaHo4 |  | Holguin | Frank Pais | 20.633719, -75.233427 | 10 |  | Burro Cemsa | Bluggoe | ABB | 2016-2018 |
| cub0097 | MosGu1 |  | Guantanamo | Baracoa | 20.261361, -74.422292 | 13 |  | Burro Cemsa | Bluggoe | ABB | 2016-2018 |
| cub0098 | MosGu2 |  | Guantanamo | Baracoa | 20.261302, -74.422075 | 13 |  | Burro Cemsa | Bluggoe | ABB | 2016-2018 |
| cub0099 | MosGu3.1 |  | Guantanamo | Baracoa | 20.261177, -74.421611 | 14 |  | Burro Cemsa | Bluggoe | ABB | 2016-2018 |
| cub0100 | MosGu3.2 |  | Guantanamo | Baracoa | 20.261177, -74.421611 | 14 |  | Burro Cemsa | Bluggoe | ABB | 2016-2018 |
| cub0101 | MosGu8 |  | Guantanamo | Baracoa | 20.261212, -74.421655 | 14 |  | Burro Cemsa | Bluggoe | ABB | 2016-2018 |
| cub0102 | MosGu4 |  | Guantanamo | Baracoa | 20.257265, -74.414429 | 12 |  | Burro Criollo | Bluggoe | ABB | 2016-2018 |
| cub0103 | MosGu5 |  | Guantanamo | Baracoa | 20.257234, -74.414065 | 12 |  | Burro Criollo | Bluggoe | ABB | 2016-2018 |
| cub0104 | MosGu6 |  | Guantanamo | Baracoa | 20.246432, -74.423753 | 12 |  | Burro CEMSA | Bluggoe | ABB | 2016-2018 |
| cub0105 | MosGu7 |  | Guantanamo | Baracoa | 20.245011, -74.426939 | 12 |  | Manzano vietnamita | Pisang Awak | ABB | 2016-2018 |
| cub0106 | SaLGu1 |  | Guantanamo | Baracoa | 20.294545, -74.436374 | 14 |  | Burro CEMSA | Bluggoe | ABB | 2016-2018 |
| cub0107 | SaLGu2 |  | Guantanamo | Baracoa | 20.294402, -74.436478 | 14 |  | Burro CEMSA | Bluggoe | ABB | 2016-2018 |

**Table S2**. (Continue)

| **DArT id** | **Isolate Code^1^** |  | **Geographical origin** | | | |  | **Host** | | | **Survey** |
| --- | --- | --- | --- | --- | --- | --- | --- | --- | --- | --- | --- |
|  |  |  | **Province** | **Municipality** | **coordinates** | **MASL^2^** |  | **Cultivar** | **Subgroup** | **genotype** |  |
| cub0108 | SaSGu1.1 |  | Guantanamo | San Antonio del Sur | 19.994528, -74.98861 | 17 |  | Burro CEMSA | Bluggoe | ABB | 2016-2018 |
| cub0109 | SaSGu1.2 |  | Guantanamo | San Antonio del Sur | 19.994528, -74.98861 | 17 |  | Burro CEMSA | Bluggoe | ABB | 2016-2018 |
| cub0110 | SaSGu2 |  | Guantanamo | San Antonio del Sur | 19.994029, -74.987754 | 17 |  | Burro CEMSA | Bluggoe | ABB | 2016-2018 |
| cub0111 | SaSGu3.1 |  | Guantanamo | San Antonio del Sur | 19.987371, -74.982173 | 14 |  | Burro CEMSA | Bluggoe | ABB | 2016-2018 |
| cub0112 | SaSGu3.2 |  | Guantanamo | San Antonio del Sur | 19.987371, -74.982173 | 14 |  | Burro CEMSA | Bluggoe | ABB | 2016-2018 |
| cub0113 | SaSGu4 |  | Guantanamo | San Antonio del Sur | 19.986947, -74.983302 | 14 |  | Burro CEMSA | Bluggoe | ABB | 2016-2018 |
| cub0114 | LaMSC1.1 |  | Santiago de Cuba | Songo La Maya | 20.167602, -75.656472 | 253 |  | Burro CEMSA | Bluggoe | ABB | 2016-2018 |
| cub0115 | LaMSC1.2 |  | Santiago de Cuba | Songo La Maya | 20.167602, -75.656472 | 253 |  | Burro CEMSA | Bluggoe | ABB | 2016-2018 |
| cub0116 | LaMSC2.1 |  | Santiago de Cuba | Songo La Maya | 20.16981, -75.684817 | 262 |  | Burro CEMSA | Bluggoe | ABB | 2016-2018 |
| cub0117 | LaMSC2.2 |  | Santiago de Cuba | Songo La Maya | 20.16981, -75.684817 | 262 |  | Burro CEMSA | Bluggoe | ABB | 2016-2018 |
| cub0118 | CriSC |  | Santiago de Cuba | Santiago de Cuba | 20.12074, -75.749968 | 200 |  | Burro CEMSA | Bluggoe | ABB | 2016-2018 |
| cub0119 | JaMay1.1 |  | Mayabeque | Jaruco | 23.10003, -82.07 | 68 |  | Manzano vietnamita | Pisang Awak | ABB | 2016-2018 |
| cub0120 | JaMay1.2 |  | Mayabeque | Jaruco | 23.10003, -82.07 | 68 |  | Manzano vietnamita | Pisang Awak | ABB | 2016-2018 |
| cub0121 | JaMay2 |  | Mayabeque | Jaruco | 23.100081, -82.069980 | 68 |  | Manzano vietnamita | Pisang Awak | ABB | 2016-2018 |
| cub0122 | JaMay3.1 |  | Mayabeque | Jaruco | 23.09985, -82.0701 | 68 |  | Manzano vietnamita | Pisang Awak | ABB | 2016-2018 |
| cub0123 | JaMay3.2 |  | Mayabeque | Jaruco | 23.09985, -82.0701 | 68 |  | Manzano vietnamita | Pisang Awak | ABB | 2016-2018 |
| cub0124 | SJMay1 |  | Mayabeque | San José | 22.931547, -82.027809 | 103 |  | Burro enano | Bluggoe | ABB | 2016-2018 |
| cub0125 | SJMay2.1 |  | Mayabeque | San José | 22.932981, -82.039293 | 103 |  | Burro enano | Bluggoe | ABB | 2016-2018 |
| cub0126 | SJMay2.2 |  | Mayabeque | San José | 22.932981, -82.039293 | 103 |  | Burro enano | Bluggoe | ABB | 2016-2018 |
| cub0128 | SJMay3.2 |  | Mayabeque | San José | 22.932295, -82.045842 | 103 |  | Burro CEMSA | Bluggoe | ABB | 2016-2018 |
| cub0129 | SJMay4 |  | Mayabeque | San José | 22.933316, -82.046872 | 103 |  | Manzano vietnamita | Pisang Awak | ABB | 2016-2018 |
| cub0130 | CabSS1 |  | Sancti Spiritus | Cabaiguan | 22.062181, -79.542205 | 152 |  | Manzano vietnamita | Pisang Awak | ABB | 2016-2018 |
| cub0131 | CabSS2 |  | Sancti Spiritus | Cabaiguan | 22.062141, -79.542425 | 152 |  | Manzano vietnamita | Pisang Awak | ABB | 2016-2018 |
| cub0132 | YagSS1 |  | Sancti Spiritus | Yaguajay | 22.263664, -79.255669 | 196 |  | Manzano vietnamita | Pisang Awak | ABB | 2016-2018 |
| cub0133 | YagSS2 |  | Sancti Spiritus | Yaguajay | 22.263654, -79.255779 | 196 |  | Manzano vietnamita | Pisang Awak | ABB | 2016-2018 |

**Table S2**. (Continue)

| **DArT id** | **Isolate Code^1^** |  | **Geographical origin** | | | |  | **Host** | | | **Survey** |
| --- | --- | --- | --- | --- | --- | --- | --- | --- | --- | --- | --- |
|  |  |  | **Province** | **Municipality** | **coordinates** | **MASL^2^** |  | **Cultivar** | **Subgroup** | **genotype** |  |
| cub0134 | YagSS3 |  | Sancti Spiritus | Yaguajay | 22.263668, -79.255651 | 196 |  | Manzano vietnamita | Pisang Awak | ABB | 2016-2018 |
| cub0135 | StCVC1 |  | Villa Clara | Santa Clara | 22.419466, -79.938231 | 114 |  | Manzano vietnamita | Pisang Awak | ABB | 2016-2018 |
| cub0136 | StCVC2 |  | Villa Clara | Santa Clara | 22.457706, -79.859251 | 99 |  | Manzano vietnamita | Pisang Awak | ABB | 2016-2018 |
| cub0137 | CaVC2 |  | Villa Clara | Caibarién | 22.502725, -79.502266 | 10 |  | Manzano vietnamita | Pisang Awak | ABB | 2016-2018 |
| cub0138 | CaVC3 |  | Villa Clara | Caibarién | 22.502241, -79.503412 | 10 |  | Manzano vietnamita | Pisang Awak | ABB | 2016-2018 |
| cub0139 | CamVC1 |  | Villa Clara | Camajuani | 22.556938, -79.722614 | 11 |  | Burro Cemsa | Bluggoe | ABB | 2016-2018 |
| cub0140 | CiVC1 |  | Villa Clara | Cifuentes | 22.565426, -80.039095 | 119 |  | Manzano vietnamita | Pisang Awak | ABB | 2016-2018 |
| cub0141 | CiVC2 |  | Villa Clara | Cifuentes | 22.565523, -80.038918 | 119 |  | Manzano vietnamita | Pisang Awak | ABB | 2016-2018 |
| cub0142 | CumCF3.1 |  | Cienfuegos | Cumanayagua | 22.139874, -80.189647 | 292 |  | Burro Cemsa | Bluggoe | ABB | 2016-2018 |
| cub0143 | CumCF3.2 |  | Cienfuegos | Cumanayagua | 22.139874, -80.189647 | 292 |  | Burro Cemsa | Bluggoe | ABB | 2016-2018 |
| cub0144 | ManVC1 |  | Villa Clara | Manicaragua | 22.068554, -79.976483 | 386 |  | Manzano vietnamita | Pisang Awak | ABB | 2016-2018 |
| cub0145 | MAnVC1-1 |  | Villa Clara | Manicaragua | 22.068554, -79.976483 | 386 |  | Manzano vietnamita | Pisang Awak | ABB | 2016-2018 |
| cub0146 | ManVC2 |  | Villa Clara | Manicaragua | 22.068520, -79.976453 | 386 |  | Manzano vietnamita | Pisang Awak | ABB | 2016-2018 |
| cub0147 | ManVC2-1 |  | Villa Clara | Manicaragua | 22.068520, -79.976453 | 386 |  | Manzano vietnamita | Pisang Awak | ABB | 2016-2018 |
| cub0149 | PalPR1 |  | Pinar del Rio | Los Palacios | 22.583139, -83.281703 | 49 |  | Burro Cemsa | Bluggoe | ABB | 2016-2018 |
| cub0150 | PalPR2 |  | Pinar del Rio | Los Palacios | 22.583167, -83.281708 | 49 |  | Burro Cemsa | Bluggoe | ABB | 2016-2018 |
| cub0151 | PalPR3 |  | Pinar del Rio | Los Palacios | 22.582946, -83.282194 | 46 |  | Burro Cemsa | Bluggoe | ABB | 2016-2018 |
| cub0152 | PalPR4 |  | Pinar del Rio | Los Palacios | 22.583670, -83.283490 | 47 |  | Burro Cemsa | Bluggoe | ABB | 2016-2018 |
| cub0153 | PalPR5 |  | Pinar del Rio | Los Palacios | 22.583811, -83.283559 | 50 |  | Burro Cemsa | Bluggoe | ABB | 2016-2018 |
| cub0154 | PalPR6 |  | Pinar del Rio | Los Palacios | 22.583817, -83.283649 | 50 |  | Burro Cemsa | Bluggoe | ABB | 2016-2018 |
| cub0155 | PalPR7 |  | Pinar del Rio | Los Palacios | 22.641865, -83.21056 | 71 |  | Manzano vietnamita | Pisang Awak | ABB | 2016-2018 |
| cub0156 | PalPR8 |  | Pinar del Rio | Los Palacios | 22.64187, -83.210544 | 71 |  | Manzano vietnamita | Pisang Awak | ABB | 2016-2018 |
| cub0157 | ViPR2 |  | Pinar del Rio | Viñales | 22.689105, -83.70567 | 63 |  | Burro enano | Bluggoe | ABB | 2016-2018 |
| cub0158 | ViPR3 |  | Pinar del Rio | Viñales | 22.689105, -83.70567 | 63 |  | Burro enano | Bluggoe | ABB | 2016-2018 |
| cub0159 | PalmPR1 |  | Pinar del Rio | La Palma | 22.730729, -83.610914 | 67 |  | Burro Cemsa | Bluggoe | ABB | 2016-2018 |

**Table S2**. (Continue)

| **DArT id** | **Isolate Code^1^** |  | **Geographical origin** | | | |  | **Host** | | | **Survey** |
| --- | --- | --- | --- | --- | --- | --- | --- | --- | --- | --- | --- |
|  |  |  | **Province** | **Municipality** | **coordinates** | **MASL^2^** |  | **Cultivar** | **Subgroup** | **genotype** |  |
| cub0160 | PalmPR2 |  | Pinar del Rio | La Palma | 22.730729, -83.610914 | 67 |  | Burro Cemsa | Bluggoe | ABB | 2016-2018 |
| cub0161 | PalmPR3.1 |  | Pinar del Rio | La Palma | 22.808993, -83.511016 | 16 |  | Burro Cemsa | Bluggoe | ABB | 2016-2018 |
| cub0162 | PalmPR3.2 |  | Pinar del Rio | La Palma | 22.808993, -83.511016 | 16 |  | Burro Cemsa | Bluggoe | ABB | 2016-2018 |
| cub0164 | GuaPR1 |  | Pinar del Rio | Guane | 22.188265, -84.205815 | 12 |  | Burro CEMSA | Bluggoe | ABB | 2016-2018 |
| cub0165 | GuaPR2 |  | Pinar del Rio | Guane | 22.188265, -84.205815 | 30 |  | Burro CEMSA | Bluggoe | ABB | 2016-2018 |
| cub0166 | SanJPR1 |  | Pinar del Rio | San Juan y Martínez | 22.228536, -83.877718 | 30 |  | Burro CEMSA | Bluggoe | ABB | 2016-2018 |
| cub0167 | SanJPR2 |  | Pinar del Rio | San Juan y Martínez | 22.65457, -83.842405 | 15 |  | Manzano vietnamita | Pisang Awak | ABB | 2016-2018 |
| cub0168 | SanJPR3 |  | Pinar del Rio | San Juan y Martínez | 22.65457, -83.842405 | 26 |  | Manzano vietnamita | Pisang Awak | ABB | 2016-2018 |
| cub0169 | PinPR |  | Pinar del Rio | Pinar de Río | 22.420475, -83.689995 | 26 |  | Burro CEMSA | Bluggoe | ABB | 2016-2018 |
| cub0170 | SanLPR1 |  | Pinar del Rio | San Luis | 22.267129, -83.769663 | 45 |  | Burro enano | Bluggoe | ABB | 2016-2018 |
| cub0171 | SanLPR2 |  | Pinar del Rio | San Luis | 22.276589, -83.75275 | 18 |  | Manzano vietnamita | Pisang Awak | ABB | 2016-2018 |
| cub0172 | SanLPR3 |  | Pinar del Rio | San Luis | 22.347291, -83.789534 | 21 |  | Burro CEMSA | Bluggoe | ABB | 2016-2018 |
| cub0173 | PerMat1 |  | Matanzas | Perico | 22.790785, -81.063242 | 30 |  | Manzano vietnamita | Pisang Awak | ABB | 2016-2018 |
| cub0174 | PerMat2.1 |  | Matanzas | Perico | 22.790563, -81.062872 | 31 |  | Manzano vietnamita | Pisang Awak | ABB | 2016-2018 |
| cub0175 | PerMat2.2 |  | Matanzas | Perico | 22.790563, -81.062872 | 31 |  | Manzano vietnamita | Pisang Awak | ABB | 2016-2018 |
| cub0176 | Art3Art1 |  | Artemisa | Artemisa | 22.795743, -82.763285 | 33 |  | Manzano vietnamita | Pisang Awak | ABB | 2016-2018 |
| cub0177 | Art3Art2 |  | Artemisa | Artemisa | 22.795743, -82.763285 | 33 |  | Manzano vietnamita | Pisang Awak | ABB | 2016-2018 |
| cub0178 | Art3Art3 |  | Artemisa | Artemisa | 22.795743, -82.763285 | 33 |  | Manzano vietnamita | Pisang Awak | ABB | 2016-2018 |
| cub0179 | ManVC3 |  | Villa Clara | Manicaragua | 22.068519, -79.976387 | 386 |  | Manzano vietnamita | Pisang Awak | ABB | 2016-2018 |
| cub0180 | R-1 |  | Villa Clara | Santo Domingo | 22.586431, -80.224707 | 48 |  | Manzano | Silk | AAB | 1996-2000 |

^1^-Isolate code in the microbial culture collection of Instituto de Investigaciones de Sanidad Vegetal (Cuba). ^2^- MASL: Meters above sea level.
