## Supplementary material for "A deep genetic analysis of banana Fusarium wilt pathogens of Cuba in a Latin American and Caribbean diversity landscape": Table S3

**Table S3**. Identification of 170 Cuban *Fusarium* isolates that cause Fusarium wilt of banana, their proposed *Fusarium* species name derived from genotyping-by-sequencing (DArTseq) data and attributed clade, vegetative compatibility group, and race identification, either by PCR or phenotyping.

| **DArT id** | **Isolate Code^1^** |  | **Origin** | | |  | **Identification** | | | | | | | **Mating type** |
| --- | --- | --- | --- | --- | --- | --- | --- | --- | --- | --- | --- | --- | --- | --- |
|  |  |  | **Province** | **Cultivar** | **Subgroup** |  | **Species^2^** | **Clade^3^** | **VCG^4^** | **Race^5^** | **Fox^a^** | **TR4^b,c^** | **R1/R2^d^** |  |
| cub0001 | IJ-1 |  | Isla de la Juventud | Burro CEMSA | Bluggoe |  | *F. tardichlamydosporum* | c2 | 0124 | R2 | + | - | + | *mat1-2* |
| cub0002 | SC-3 |  | Santiago de Cuba | Burro criollo | Bluggoe |  | *F. tardichlamydosporum* | c2 | 0128 | R2 | + | - | + | *mat1-2* |
| cub0003 | C.Esm1.3 |  | Camaguey | Burro criollo | Bluggoe |  | *F. tardichlamydosporum* | c2 | 0124/5 | R2 | + | - | + | *mat1-2* |
| cub0004 | Cca1.1 |  | Camaguey | Manzano | Silk |  | *F. tardichlamydosporum* | c2 | 01210 | R1 | + | - | + | *mat1-2* |
| cub0005 | R.Go |  | Havana | Manzano | Silk |  | *F. tardichlamydosporum* | c2 | 01210 | R1 | + | - | + | *mat1-2* |
| cub0006 | Cub9 |  | Villa Clara | Gros Michel | Gros Michel |  | *F. purpurascens* | c1 | 01210 | n.d. | + | - | + | *mat1-1* |
| cub0007 | M3 |  | Villa Clara | Manzano | Silk |  | *F. purpurascens* | c1 | 01210 | n.d. | + | - | + | *mat1-2* |
| cub0008 | C.Esm1.1 |  | Camaguey | Burro criollo | Bluggoe |  | *F. tardichlamydosporum* | c2 | 0124/5 | R2 | + | - | + | *mat1-2* |
| cub0009 | CGua 1.1 |  | Camaguey | Burro criollo | Bluggoe |  | *F. tardicrescens* | c2 | 01210 | R2 | + | - | + | *mat1-1* |
| cub0010 | C.V 2.1 |  | Camaguey | Manzano | Silk |  | *F. purpurascens* | c1 | 01210 | R1 | + | - | + | *mat1-1* |
| cub0011 | ESB-1 |  | Camaguey | Burro criollo | Bluggoe |  | *F. tardichlamydosporum* | c2 | 01210 | R2 | + | - | + | *mat1-2* |
| cub0012 | Cam3 |  | Camaguey | Manzano | Silk |  | *F. purpurascens* | c1 | 01210 | R1* | + | - | + | *mat1-1* |
| cub0013 | SC-2 |  | Santiago de Cuba | FHIA-03 | FHIA hybrid |  | *F. tardichlamydosporum* | c2 | 01210 | n.d. | + | - | + | *mat1-2* |
| cub0014 | GuCUM |  | Guantanamo | Burro CEMSA | Bluggoe |  | *F. tardichlamydosporum* | c2 | 0124 | R2 | + | - | + | *mat1-2* |
| cub0015 | GuBaCu |  | Guantanamo | Burro Criollo | Bluggoe |  | *F. tardichlamydosporum* | c2 | 0124 | R2 | + | - | + | *mat1-2* |
| cub0016 | GuMa |  | Guantanamo | Burro Criollo | Bluggoe |  | *F. tardichlamydosporum* | c2 | 0124 | R2 | + | - | + | *mat1-2* |
| cub0017 | TUPP-1 |  | Las Tunas | Manzano | Silk |  | *F. tardichlamydosporum* | c2 | 01210 | R1 | + | - | + | *mat1-1* |
| cub0018 | GuBaPo |  | Guantanamo | Burro Criollo | Bluggoe |  | *F. tardichlamydosporum* | c2 | 0124 | R2 | + | - | + | *mat1-2* |
| cub0020 | MoGu 1 |  | Artemisa | Manzano vietnamita | Pisang Awak |  | *F. tardichlamydosporum* | c2 | 0124 | R2 | + | - | + | *mat1-2* |
| cub0021 | MoGu 3 |  | Artemisa | Manzano vietnamita | Pisang Awak |  | *F. tardichlamydosporum* | c2 | 0124 | R2 | + | - | + | *mat1-2* |
| cub0022 | WaHa |  | La Habana | Burro CEMSA | Bluggoe |  | *F. phialophorum* | c2 | n.d. | R2* | + | - | + | *mat1-2* |
| cub0023 | PaGu |  | Artemisa | Manzano vietnamita | Pisang Awak |  | *F. tardichlamydosporum* | c2 | 0124 | R2 | + | - | + | *mat1-2* |
| cub0024 | AlqHa 1 |  | Artemisa | Manzano vietnamita | Pisang Awak |  | *F. tardichlamydosporum* | c2 | n.d. | n.d. | + | - | + | *mat1-2* |
| cub0025 | GuHa |  | Artemisa | Manzano vietnamita | Pisang Awak |  | *F. tardichlamydosporum* | c2 | n.d. | n.d. | + | - | + | *mat1-2* |
| cub0026 | PeBa |  | Guantanamo | Burro CEMSA | Bluggoe |  | *F. tardichlamydosporum* | c2 | 0124 | n.d. | + | - | + | *mat1-2* |

**Table S3**. (Continue)

| **DArT id** | **Isolate Code^1^** |  | **Origin** | | |  | **Identification** | | | | | | | **Mating type** |
| --- | --- | --- | --- | --- | --- | --- | --- | --- | --- | --- | --- | --- | --- | --- |
|  |  |  | **Province** | **Cultivar** | **Subgroup** |  | **Species^2^** | **Clade^3^** | **VCG^4^** | **Race^5^** | **Fox^a^** | **TR4^b,c^** | **R1/R2^d^** |  |
| cub0027 | ToaGu |  | Guantanamo | Burro criollo | Bluggoe |  | *F. tardichlamydosporum* | c2 | 0124 | R2 | + | - | + | *mat1-2* |
| cub0028 | Bar 1 |  | Guantanamo | Burro criollo | Bluggoe |  | *F. tardichlamydosporum* | c2 | 0124 | R2 | + | - | + | *mat1-2* |
| cub0029 | MaHo 1 |  | Holguin | Burro CEMSA | Bluggoe |  | *F. tardichlamydosporum* | c2 | 0124 | R2 | + | - | + | *mat1-2* |
| cub0030 | BaHo 1 |  | Holguin | Burro criollo | Bluggoe |  | *F. tardichlamydosporum* | c2 | 0124 | R2 | + | - | + | *mat1-2* |
| cub0031 | BaHo 2 |  | Holguin | Burro criollo | Bluggoe |  | *F. tardichlamydosporum* | c2 | 0124 | R2 | + | - | + | *mat1-2* |
| cub0032 | SaHo |  | Holguin | Burro CEMSA | Bluggoe |  | *F. tardichlamydosporum* | c2 | 0124 | R2 | + | - | + | *mat1-2* |
| cub0033 | Bar 3 |  | Guantanamo | Manzano | Silk |  | *F. tardichlamydosporum* | c2 | 01210 | R1 | + | - | + | *mat1-2* |
| cub0034 | R-2 |  | Villa Clara | Burro criollo | Bluggoe |  | *F. tardicrescens* | c2 | 01210 | R2* | + | - | + | *mat1-2* |
| cub0035 | JCien |  | Cienfuegos | Manzano vietnamita | Pisang Awak |  | *F. tardichlamydosporum* | c2 | n.d. | R1* | + | - | + | *mat1-1* |
| cub0037 | CaVC |  | Villa Clara | Manzano vietnamita | Pisang Awak |  | *F. tardichlamydosporum* | c2 | n.d. | n.d. | + | - | + | *mat1-2* |
| cub0038 | ToBaGu |  | Guantanamo | Burro CEMSA | Bluggoe |  | *F. tardichlamydosporum* | c2 | n.d. | n.d. | + | - | + | *mat1-2* |
| cub0039 | NarBaGu |  | Guantanamo | Burro CEMSA | Bluggoe |  | *F. tardichlamydosporum* | c2 | n.d. | n.d. | + | - | + | *mat1-2* |
| cub0040 | PGGua1 |  | Santiago de Cuba | Burro Criollo | Bluggoe |  | *F. tardichlamydosporum* | c2 | n.d. | n.d. | + | - | + | *mat1-2* |
| cub0041 | PGGua2 |  | Santiago de Cuba | Manzano vietnamita | Pisang Awak |  | *F. tardichlamydosporum* | c2 | n.d. | n.d. | + | - | + | *mat1-2* |
| cub0042 | BaGra |  | Granma | Burro CEMSA | Bluggoe |  | *F. tardichlamydosporum* | c2 | n.d. | n.d. | + | - | + | *mat1-2* |
| cub0043 | PunBra1 |  | La Habana | Manzano vietnamita | Pisang Awak |  | *F. tardichlamydosporum* | c2 | n.d. | R1* | + | - | + | *mat1-2* |
| cub0044 | PunBra2 |  | La Habana | Manzano vietnamita | Pisang Awak |  | *F. tardichlamydosporum* | c2 | n.d. | n.d. | + | - | + | *mat1-2* |
| cub0045 | BatMay1 |  | Mayabeque | Burro enano | Bluggoe |  | *F. tardichlamydosporum* | c2 | n.d. | n.d. | + | - | + | *mat1-2* |
| cub0046 | BatMay2 |  | Mayabeque | Manzano vietnamita | Pisang Awak |  | *F. tardichlamydosporum* | c2 | n.d. | n.d. | + | - | + | *mat1-2* |
| cub0047 | Min1Art |  | Artemisa | Manzano vietnamita | Pisang Awak |  | *F. tardichlamydosporum* | c2 | n.d. | n.d. | + | - | + | *mat1-2* |
| cub0048 | MaySC1 |  | Santiago de Cuba | Burro Criollo | Bluggoe |  | *F. tardichlamydosporum* | c2 | n.d. | n.d. | + | - | + | *mat1-2* |
| cub0049 | MaySC2 |  | Santiago de Cuba | Burro Criollo | Bluggoe |  | *F. tardichlamydosporum* | c2 | n.d. | n.d. | + | - | + | *mat1-2* |
| cub0050 | CumCF1 |  | Cienfuegos | Burro CEMSA | Bluggoe |  | *F. tardichlamydosporum* | c2 | n.d. | n.d. | + | - | + | *mat1-2* |
| cub0051 | CumCF2 |  | Cienfuegos | Burro CEMSA | Bluggoe |  | *F. tardichlamydosporum* | c2 | n.d. | n.d. | + | - | + | *mat1-2* |
| cub0052 | BoyHa |  | La Habana | Manzano vietnamita | Pisang Awak |  | *F. tardichlamydosporum* | c2 | n.d. | n.d. | + | - | + | *mat1-2* |
| cub0053 | CotHa |  | La Habana | Manzano vietnamita | Pisang Awak |  | *F. tardichlamydosporum* | c2 | n.d. | n.d. | + | - | + | *mat1-2* |

**Table S3**. (Continue)

| **DArT id** | **Isolate Code^1^** |  | **Origin** | | |  | **Identification** | | | | | | | **Mating type** |
| --- | --- | --- | --- | --- | --- | --- | --- | --- | --- | --- | --- | --- | --- | --- |
|  |  |  | **Province** | **Cultivar** | **Subgroup** |  | **Species^2^** | **Clade^3^** | **VCG^4^** | **Race^5^** | **Fox^a^** | **TR4^b,c^** | **R1/R2^d^** |  |
| cub0054 | GuaHa1 |  | La Habana | Manzano vietnamita | Pisang Awak |  | *F. tardichlamydosporum* | c2 | n.d. | n.d. | + | - | + | *mat1-2* |
| cub0055 | C.Esm1.2 |  | Camaguey | Burro CEMSA | Bluggoe |  | *F. tardichlamydosporum* | c2 | 0124/5 | R2 | + | - | + | *mat1-2* |
| cub0056 | C.Esm2.1 |  | Camaguey | Burro criollo | Bluggoe |  | *F. tardichlamydosporum* | c2 | 0124/5 | R2 | + | - | + | *mat1-2* |
| cub0057 | ESB-5 |  | Camaguey | Burro criollo | Bluggoe |  | *F. tardichlamydosporum* | c2 | 0124 | R2 | + | - | + | *mat1-2* |
| cub0058 | SCGu |  | Santiago de Cuba | Burro CEMSA | Bluggoe |  | *F. tardichlamydosporum* | c2 | 0124 | R2 | + | - | + | *mat1-2* |
| cub0059 | Bar 2 |  | Guantanamo | Manzano | Silk |  | *F. tardichlamydosporum* | c2 | 0124 | R2 | + | - | + | *mat1-2* |
| cub0060 | ColCA |  | Ciego de Avila | Burro criollo | Bluggoe |  | *F. tardichlamydosporum* | c2 | n.d. | n.d. | + | - | + | *mat1-2* |
| cub0061 | Min2Art |  | Artemisa | Manzano vietnamita | Pisang Awak |  | *F. tardichlamydosporum* | c2 | n.d. | n.d. | + | - | + | *mat1-2* |
| cub0063 | Art1Art-1 |  | Artemisa | Manzano vietnamita | Pisang Awak |  | *F. tardichlamydosporum* | c2 | n.d. | n.d. | + | - | + | *mat1-2* |
| cub0064 | Art1Art-2 |  | Artemisa | Burro CEMSA | Bluggoe |  | *F. tardichlamydosporum* | c2 | n.d. | n.d. | + | - | + | *mat1-2* |
| cub0065 | Art2Art-1 |  | Artemisa | Burro CEMSA | Bluggoe |  | *F. tardichlamydosporum* | c2 | n.d. | n.d. | + | - | + | *mat1-2* |
| cub0067 | MelArt1.1 |  | Mayabeque | Burro Criollo | Bluggoe |  | *F. tardichlamydosporum* | c2 | n.d. | n.d. | + | - | + | *mat1-2* |
| cub0068 | MelArt1.2 |  | Mayabeque | Manzano vietnamita | Pisang Awak |  | *F. tardichlamydosporum* | c2 | n.d. | n.d. | + | - | + | *mat1-2* |
| cub0069 | Front1Art1 |  | Artemisa | Burro CEMSA | Bluggoe |  | *F. tardichlamydosporum* | c2 | n.d. | n.d. | + | - | + | *mat1-2* |
| cub0070 | Front1Art2 |  | Artemisa | Manzano vietnamita | Pisang Awak |  | *F. tardichlamydosporum* | c2 | n.d. | n.d. | + | - | + | *mat1-2* |
| cub0071 | Front2Art |  | Artemisa | Manzano vietnamita | Pisang Awak |  | *F. tardichlamydosporum* | c2 | n.d. | n.d. | + | - | + | *mat1-1* |
| cub0072 | FloCam1 |  | Camaguey | Burro enano | Bluggoe |  | *F. tardichlamydosporum* | c2 | n.d. | n.d. | + | - | + | *mat1-2* |
| cub0073 | CesCam1 |  | Camaguey | Manzano vietnamita | Pisang Awak |  | *F. tardichlamydosporum* | c2 | n.d. | n.d. | + | - | + | *mat1-2* |
| cub0075 | LisHa2 |  | La Habana | Manzano vietnamita | Pisang Awak |  | *F. tardichlamydosporum* | c2 | n.d. | n.d. | + | - | + | *mat1-2* |
| cub0076 | TolHa |  | La Habana | Burro Criollo | Bluggoe |  | *F. tardichlamydosporum* | c2 | n.d. | n.d. | + | - | + | *mat1-2* |
| cub0077 | AldHa |  | La Habana | Burro Criollo | Bluggoe |  | *F. tardichlamydosporum* | c2 | n.d. | n.d. | + | - | + | *mat1-2* |
| cub0078 | FloCam2 |  | Camaguey | Burro CEMSA | Bluggoe |  | *F. tardichlamydosporum* | c2 | n.d. | n.d. | + | - | + | *mat1-2* |
| cub0079 | FloCam3 |  | Camaguey | Burro CEMSA | Bluggoe |  | *F. tardichlamydosporum* | c2 | n.d. | n.d. | + | - | + | *mat1-2* |
| cub0080 | FloCam4 |  | Camaguey | Manzano vietnamita | Pisang Awak |  | *F. tardichlamydosporum* | c2 | n.d. | n.d. | + | - | + | *mat1-1* |

**Table S3**. (Continue)

| **DArT id** | **Isolate Code^1^** |  | **Origin** | | |  | **Identification** | | | | | | | **Mating type** |
| --- | --- | --- | --- | --- | --- | --- | --- | --- | --- | --- | --- | --- | --- | --- |
|  |  |  | **Province** | **Cultivar** | **Subgroup** |  | **Species^2^** | **Clade^3^** | **VCG^4^** | **Race^5^** | **Fox^a^** | **TR4^b,c^** | **R1/R2^d^** |  |
| cub0081 | CCam1 |  | Camaguey | Manzano vietnamita | Pisang Awak |  | *F. tardichlamydosporum* | c2 | n.d. | n.d. | + | - | + | *mat1-2* |
| cub0082 | CCam2 |  | Camaguey | Burro enano | Bluggoe |  | *F. tardichlamydosporum* | c2 | n.d. | n.d. | + | - | + | *mat1-2* |
| cub0084 | CCam3.2 |  | Camaguey | Burro enano | Bluggoe |  | *F. tardichlamydosporum* | c2 | n.d. | n.d. | + | - | + | *mat1-2* |
| cub0085 | MaHo2 |  | Holguin | Burro Cemsa | Bluggoe |  | *F. tardichlamydosporum* | c2 | n.d. | n.d. | + | - | + | *mat1-2* |
| cub0086 | MaHo3 |  | Holguin | Burro Cemsa | Bluggoe |  | *F. tardichlamydosporum* | c2 | n.d. | n.d. | + | - | + | *mat1-2* |
| cub0087 | MaHo4 |  | Holguin | Burro Cemsa | Bluggoe |  | *F. tardichlamydosporum* | c2 | n.d. | n.d. | + | - | + | *mat1-2* |
| cub0088 | MaHo5 |  | Holguin | Burro Cemsa | Bluggoe |  | *F. tardichlamydosporum* | c2 | n.d. | R2* | + | - | + | *mat1-2* |
| cub0089 | MaHo6 |  | Holguin | Burro Cemsa | Bluggoe |  | *F. tardichlamydosporum* | c2 | n.d. | n.d. | + | - | + | *mat1-2* |
| cub0090 | MaHo7 |  | Holguin | Burro Cemsa | Bluggoe |  | *F. tardichlamydosporum* | c2 | n.d. | n.d. | + | - | + | *mat1-2* |
| cub0091 | PaHo1 |  | Holguin | Burro Cemsa | Bluggoe |  | *F. tardichlamydosporum* | c2 | n.d. | n.d. | + | - | + | *mat1-2* |
| cub0092 | PaHo2 |  | Holguin | Burro Cemsa | Bluggoe |  | *F. tardichlamydosporum* | c2 | n.d. | n.d. | + | - | + | *mat1-2* |
| cub0093 | SaHo2 |  | Holguin | Burro Criollo | Bluggoe |  | *F. tardichlamydosporum* | c2 | n.d. | n.d. | + | - | + | *mat1-2* |
| cub0095 | SaHo3.2 |  | Holguin | Burro Cemsa | Bluggoe |  | *F. tardichlamydosporum* | c2 | n.d. | n.d. | + | - | + | *mat1-2* |
| cub0096 | SaHo4 |  | Holguin | Burro Cemsa | Bluggoe |  | *F. tardichlamydosporum* | c2 | n.d. | n.d. | + | - | + | *mat1-1* |
| cub0097 | MosGu1 |  | Guantanamo | Burro Cemsa | Bluggoe |  | *F. tardichlamydosporum* | c2 | n.d. | n.d. | + | - | + | *mat1-2* |
| cub0098 | MosGu2 |  | Guantanamo | Burro Cemsa | Bluggoe |  | *F. tardichlamydosporum* | c2 | n.d. | n.d. | + | - | + | *mat1-2* |
| cub0099 | MosGu3.1 |  | Guantanamo | Burro Cemsa | Bluggoe |  | *F. tardichlamydosporum* | c2 | n.d. | n.d. | + | - | + | *mat1-2* |
| cub0100 | MosGu3.2 |  | Guantanamo | Burro Cemsa | Bluggoe |  | *F. tardichlamydosporum* | c2 | n.d. | n.d. | + | - | + | *mat1-2* |
| cub0101 | MosGu8 |  | Guantanamo | Burro Cemsa | Bluggoe |  | *F. tardichlamydosporum* | c2 | n.d. | n.d. | + | - | + | *mat1-2* |
| cub0102 | MosGu4 |  | Guantanamo | Burro Criollo | Bluggoe |  | *F. tardichlamydosporum* | c2 | n.d. | n.d. | + | - | + | *mat1-2* |
| cub0103 | MosGu5 |  | Guantanamo | Burro Criollo | Bluggoe |  | *F. tardichlamydosporum* | c2 | n.d. | n.d. | + | - | + | *mat1-2* |
| cub0104 | MosGu6 |  | Guantanamo | Burro CEMSA | Bluggoe |  | *F. tardichlamydosporum* | c2 | n.d. | n.d. | + | - | + | *mat1-2* |
| cub0105 | MosGu7 |  | Guantanamo | Manzano vietnamita | Pisang Awak |  | *F. tardichlamydosporum* | c2 | n.d. | n.d. | + | - | + | *mat1-2* |
| cub0106 | SaLGu1 |  | Guantanamo | Burro CEMSA | Bluggoe |  | *F. tardichlamydosporum* | c2 | n.d. | n.d. | + | - | + | *mat1-2* |
| cub0107 | SaLGu2 |  | Guantanamo | Burro CEMSA | Bluggoe |  | *F. tardichlamydosporum* | c2 | n.d. | n.d. | + | - | + | *mat1-2* |
| cub0108 | SaSGu1.1 |  | Guantanamo | Burro CEMSA | Bluggoe |  | *F. tardichlamydosporum* | c2 | n.d. | n.d. | + | - | + | *mat1-2* |

**Table S3**. (Continue)

| **DArT id** | **Isolate Code^1^** |  | **Origin** | | |  | **Identification** | | | | | | | **Mating type** |
| --- | --- | --- | --- | --- | --- | --- | --- | --- | --- | --- | --- | --- | --- | --- |
|  |  |  | **Province** | **Cultivar** | **Subgroup** |  | **Species^2^** | **Clade^3^** | **VCG^4^** | **Race^5^** | **Fox^a^** | **TR4^b,c^** | **R1/R2^d^** |  |
| cub0109 | SaSGu1.2 |  | Guantanamo | Burro CEMSA | Bluggoe |  | *F. tardichlamydosporum* | c2 | n.d. | n.d. | + | - | + | *mat1-2* |
| cub0110 | SaSGu2 |  | Guantanamo | Burro CEMSA | Bluggoe |  | *F. tardichlamydosporum* | c2 | n.d. | n.d. | + | - | + | *mat1-2* |
| cub0111 | SaSGu3.1 |  | Guantanamo | Burro CEMSA | Bluggoe |  | *F. tardichlamydosporum* | c2 | n.d. | n.d. | + | - | + | *mat1-2* |
| cub0112 | SaSGu3.2 |  | Guantanamo | Burro CEMSA | Bluggoe |  | *F. tardichlamydosporum* | c2 | n.d. | n.d. | + | - | + | *mat1-2* |
| cub0113 | SaSGu4 |  | Guantanamo | Burro CEMSA | Bluggoe |  | *F. tardichlamydosporum* | c2 | n.d. | n.d. | + | - | + | *mat1-2* |
| cub0114 | LaMSC1.1 |  | Santiago de Cuba | Burro CEMSA | Bluggoe |  | *F. tardichlamydosporum* | c2 | n.d. | n.d. | + | - | + | *mat1-2* |
| cub0115 | LaMSC1.2 |  | Santiago de Cuba | Burro CEMSA | Bluggoe |  | *F. tardichlamydosporum* | c2 | n.d. | n.d. | + | - | + | *mat1-2* |
| cub0116 | LaMSC2.1 |  | Santiago de Cuba | Burro CEMSA | Bluggoe |  | *F. tardichlamydosporum* | c2 | n.d. | n.d. | + | - | + | *mat1-2* |
| cub0117 | LaMSC2.2 |  | Santiago de Cuba | Burro CEMSA | Bluggoe |  | *F. tardichlamydosporum* | c2 | n.d. | n.d. | + | - | + | *mat1-2* |
| cub0118 | CriSC |  | Santiago de Cuba | Burro CEMSA | Bluggoe |  | *F. tardichlamydosporum* | c2 | n.d. | n.d. | + | - | + | *mat1-2* |
| cub0119 | JaMay1.1 |  | Mayabeque | Manzano vietnamita | Pisang Awak |  | *F. tardichlamydosporum* | c2 | n.d. | n.d. | + | - | + | *mat1-2* |
| cub0120 | JaMay1.2 |  | Mayabeque | Manzano vietnamita | Pisang Awak |  | *F. tardichlamydosporum* | c2 | n.d. | n.d. | + | - | + | *mat1-2* |
| cub0121 | JaMay2 |  | Mayabeque | Manzano vietnamita | Pisang Awak |  | *F. tardichlamydosporum* | c2 | n.d. | n.d. | + | - | + | *mat1-2* |
| cub0122 | JaMay3.1 |  | Mayabeque | Manzano vietnamita | Pisang Awak |  | *F. tardichlamydosporum* | c2 | n.d. | n.d. | + | - | + | *mat1-2* |
| cub0123 | JaMay3.2 |  | Mayabeque | Manzano vietnamita | Pisang Awak |  | *F. tardichlamydosporum* | c2 | n.d. | n.d. | + | - | + | *mat1-2* |
| cub0124 | SJMay1 |  | Mayabeque | Burro enano | Bluggoe |  | *F. tardichlamydosporum* | c2 | n.d. | n.d. | + | - | + | *mat1-2* |
| cub0125 | SJMay2.1 |  | Mayabeque | Burro enano | Bluggoe |  | *F. tardichlamydosporum* | c2 | n.d. | n.d. | + | - | + | *mat1-2* |
| cub0126 | SJMay2.2 |  | Mayabeque | Burro enano | Bluggoe |  | *F. tardichlamydosporum* | c2 | n.d. | n.d. | + | - | + | *mat1-2* |
| cub0128 | SJMay3.2 |  | Mayabeque | Burro CEMSA | Bluggoe |  | *F. tardichlamydosporum* | c2 | n.d. | n.d. | + | - | + | *mat1-2* |
| cub0129 | SJMay4 |  | Mayabeque | Manzano vietnamita | Pisang Awak |  | *F. purpurascens* | c1 | n.d. | R1* | + | - | + | *mat1-1* |
| cub0130 | CabSS1 |  | Sancti Spiritus | Manzano vietnamita | Pisang Awak |  | *F. purpurascens* | c1 | n.d. | R1* | + | - | + | *mat1-1* |
| cub0131 | CabSS2 |  | Sancti Spiritus | Manzano vietnamita | Pisang Awak |  | *F. purpurascens* | c1 | n.d. | n.d. | + | - | + | *mat1-1* |
| cub0132 | YagSS1 |  | Sancti Spiritus | Manzano vietnamita | Pisang Awak |  | *F. tardichlamydosporum* | c2 | n.d. | n.d. | + | - | + | *mat1-1* |
| cub0133 | YagSS2 |  | Sancti Spiritus | Manzano vietnamita | Pisang Awak |  | *F. tardichlamydosporum* | c2 | n.d. | n.d. | + | - | + | *mat1-2* |
| cub0134 | YagSS3 |  | Sancti Spiritus | Manzano vietnamita | Pisang Awak |  | *F. tardichlamydosporum* | c2 | n.d. | n.d. | + | - | + | *mat1-2* |
| cub0135 | StCVC1 |  | Villa Clara | Manzano vietnamita | Pisang Awak |  | *F. tardichlamydosporum* | c2 | n.d. | n.d. | + | - | + | *mat1-2* |

**Table S3**. (Continue)

| **DArT id** | **Isolate Code^1^** | |  | | **Origin** | | | | | |  | | **Identification** | | | | | | | | | | | | | | **Mating type** |
| --- | --- | --- | --- | --- | --- | --- | --- | --- | --- | --- | --- | --- | --- | --- | --- | --- | --- | --- | --- | --- | --- | --- | --- | --- | --- | --- | --- |
|  |  |  |  | | **Province** | | **Cultivar** | | **Subgroup** | |  | | **Species^2^** | | **Clade^3^** | | **VCG^4^** | | **Race^5^** | | **Fox^a^** | | **TR4^b,c^** | | **R1/R2^d^** | |  |
| cub0136 | StCVC2 |  | | Villa Clara | | Manzano vietnamita | | Pisang Awak | |  | | *F. tardichlamydosporum* | | c2 | | n.d. | | n.d. | | + | | - | | + | | *mat1-2* | |
| cub0137 | CaVC2 |  | | Villa Clara | | Manzano vietnamita | | Pisang Awak | |  | | *F. tardichlamydosporum* | | c2 | | n.d. | | n.d. | | + | | - | | + | | *mat1-2* | |
| cub0138 | CaVC3 |  | | Villa Clara | | Manzano vietnamita | | Pisang Awak | |  | | *F. tardichlamydosporum* | | c2 | | n.d. | | n.d. | | + | | - | | + | | *mat1-2* | |
| cub0139 | CamVC1 | |  | | Villa Clara | | Burro Cemsa | | Bluggoe | |  | | *F. tardichlamydosporum* | | c2 | | n.d. | | n.d. | | + | | - | | + | | *mat1-2* |
| cub0140 | CiVC1 | |  | | Villa Clara | | Manzano vietnamita | | Pisang Awak | |  | | *F. purpurascens* | | c1 | | n.d. | | R1* | | + | | - | | + | | *mat1-1* |
| cub0141 | CiVC2 | |  | | Villa Clara | | Manzano vietnamita | | Pisang Awak | |  | | *F. purpurascens* | | c1 | | n.d. | | n.d. | | + | | - | | + | | *mat1-1* |
| cub0142 | CumCF3.1 | |  | | Cienfuegos | | Burro Cemsa | | Bluggoe | |  | | *F. tardichlamydosporum* | | c2 | | n.d. | | n.d. | | + | | - | | + | | *mat1-2* |
| cub0143 | CumCF3.2 | |  | | Cienfuegos | | Burro Cemsa | | Bluggoe | |  | | *F. tardichlamydosporum* | | c2 | | n.d. | | n.d. | | + | | - | | + | | *mat1-2* |
| cub0144 | ManVC1 | |  | | Villa Clara | | Manzano vietnamita | | Pisang Awak | |  | | *F. tardichlamydosporum* | | c2 | | n.d. | | n.d. | | + | | - | | + | | *mat1-2* |
| cub0145 | MAnVC1-1 | |  | | Villa Clara | | Manzano vietnamita | | Pisang Awak | |  | | *F. tardichlamydosporum* | | c2 | | n.d. | | R1* | | + | | - | | + | | *mat1-2* |
| cub0146 | ManVC2 | |  | | Villa Clara | | Manzano vietnamita | | Pisang Awak | |  | | *F. tardichlamydosporum* | | c2 | | n.d. | | n.d. | | + | | - | | + | | *mat1-2* |
| cub0147 | ManVC2-1 | |  | | Villa Clara | | Manzano vietnamita | | Pisang Awak | |  | | *F. tardichlamydosporum* | | c2 | | n.d. | | n.d. | | + | | - | | + | | *mat1-2* |
| cub0149 | PalPR1 | |  | | Pinar del Rio | | Burro Cemsa | | Bluggoe | |  | | *F. tardichlamydosporum* | | c2 | | n.d. | | n.d. | | + | | - | | + | | *mat1-2* |
| cub0150 | PalPR2 | |  | | Pinar del Rio | | Burro Cemsa | | Bluggoe | |  | | *F. tardichlamydosporum* | | c2 | | n.d. | | R2* | | + | | - | | + | | *mat1-2* |
| cub0151 | PalPR3 | |  | | Pinar del Rio | | Burro Cemsa | | Bluggoe | |  | | *F. tardichlamydosporum* | | c2 | | n.d. | | n.d. | | + | | - | | + | | *mat1-2* |
| cub0152 | PalPR4 | |  | | Pinar del Rio | | Burro Cemsa | | Bluggoe | |  | | *F. tardichlamydosporum* | | c2 | | n.d. | | n.d. | | + | | - | | + | | *mat1-2* |
| cub0153 | PalPR5 | |  | | Pinar del Rio | | Burro Cemsa | | Bluggoe | |  | | *F. tardichlamydosporum* | | c2 | | n.d. | | n.d. | | + | | - | | + | | *mat1-2* |
| cub0154 | PalPR6 | |  | | Pinar del Rio | | Burro Cemsa | | Bluggoe | |  | | *F. tardichlamydosporum* | | c2 | | n.d. | | n.d. | | + | | - | | + | | *mat1-2* |
| cub0155 | PalPR7 | |  | | Pinar del Rio | | Manzano vietnamita | | Pisang Awak | |  | | *F. tardichlamydosporum* | | c2 | | n.d. | | n.d. | | + | | - | | + | | *mat1-2* |
| cub0156 | PalPR8 | |  | | Pinar del Rio | | Manzano vietnamita | | Pisang Awak | |  | | *F. tardichlamydosporum* | | c2 | | n.d. | | n.d. | | + | | - | | + | | *mat1-2* |
| cub0157 | ViPR2 | |  | | Pinar del Rio | | Burro enano | | Bluggoe | |  | | *F. tardichlamydosporum* | | c2 | | n.d. | | n.d. | | + | | - | | + | | *mat1-2* |
| cub0158 | ViPR3 | |  | | Pinar del Rio | | Burro enano | | Bluggoe | |  | | *F. tardichlamydosporum* | | c2 | | n.d. | | n.d. | | + | | - | | + | | *mat1-2* |
| cub0159 | PalmPR1 | |  | | Pinar del Rio | | Burro Cemsa | | Bluggoe | |  | | *F. tardichlamydosporum* | | c2 | | n.d. | | n.d. | | + | | - | | + | | *mat1-2* |
| cub0160 | PalmPR2 | |  | | Pinar del Rio | | Burro Cemsa | | Bluggoe | |  | | *F. tardichlamydosporum* | | c2 | | n.d. | | n.d. | | + | | - | | + | | *mat1-2* |
| cub0162 | PalmPR3.2 | |  | | Pinar del Rio | | Burro Cemsa | | Bluggoe | |  | | *F. tardichlamydosporum* | | c2 | | n.d. | | n.d. | | + | | - | | + | | *mat1-2* |

**Table S3**. (Continue)

| **DArT id** | **Isolate Code^1^** |  | **Origin** | | |  | **Identification** | | | | | | | **Mating type** |
| --- | --- | --- | --- | --- | --- | --- | --- | --- | --- | --- | --- | --- | --- | --- |
|  |  |  | **Province** | **Cultivar** | **Subgroup** |  | **Species^2^** | **Clade^3^** | **VCG^4^** | **Race^5^** | **Fox^a^** | **TR4^b,c^** | **R1/R2^d^** |  |
| cub0164 | GuaPR1 |  | Pinar del Rio | Burro CEMSA | Bluggoe |  | *F. tardichlamydosporum* | c2 | n.d. | n.d. | + | - | + | *mat1-2* |
| cub0165 | GuaPR2 |  | Pinar del Rio | Burro CEMSA | Bluggoe |  | *F. tardichlamydosporum* | c2 | n.d. | n.d. | + | - | + | *mat1-2* |
| cub0166 | SanJPR1 |  | Pinar del Rio | Burro CEMSA | Bluggoe |  | *F. tardichlamydosporum* | c2 | n.d. | n.d. | + | - | + | *mat1-2* |
| cub0167 | SanJPR2 |  | Pinar del Rio | Manzano vietnamita | Pisang Awak |  | *F. tardichlamydosporum* | c2 | n.d. | n.d. | + | - | + | *mat1-2* |
| cub0168 | SanJPR3 |  | Pinar del Rio | Manzano vietnamita | Pisang Awak |  | *F. tardichlamydosporum* | c2 | n.d. | n.d. | + | - | + | *mat1-2* |
| cub0169 | PinPR |  | Pinar del Rio | Burro CEMSA | Bluggoe |  | *F. tardichlamydosporum* | c2 | n.d. | n.d. | + | - | + | *mat1-2* |
| cub0170 | SanLPR1 |  | Pinar del Rio | Burro enano | Bluggoe |  | *F. tardichlamydosporum* | c2 | n.d. | n.d. | + | - | + | *mat1-2* |
| cub0171 | SanLPR2 |  | Pinar del Rio | Manzano vietnamita | Pisang Awak |  | *F. tardichlamydosporum* | c2 | n.d. | n.d. | + | - | + | *mat1-2* |
| cub0172 | SanLPR3 |  | Pinar del Rio | Burro CEMSA | Bluggoe |  | *F. tardichlamydosporum* | c2 | n.d. | n.d. | + | - | + | *mat1-2* |
| cub0173 | PerMat1 |  | Matanzas | Manzano vietnamita | Pisang Awak |  | *F. tardichlamydosporum* | c2 | n.d. | n.d. | + | - | + | *mat1-2* |
| cub0174 | PerMat2.1 |  | Matanzas | Manzano vietnamita | Pisang Awak |  | *F. tardichlamydosporum* | c2 | n.d. | n.d. | + | - | + | *mat1-2* |
| cub0175 | PerMat2.2 |  | Matanzas | Manzano vietnamita | Pisang Awak |  | *F. tardichlamydosporum* | c2 | n.d. | n.d. | + | - | + | *mat1-2* |
| cub0176 | Art3Art1 |  | Artemisa | Manzano vietnamita | Pisang Awak |  | *F. tardichlamydosporum* | c2 | n.d. | n.d. | + | - | + | *mat1-2* |
| cub0177 | Art3Art2 |  | Artemisa | Manzano vietnamita | Pisang Awak |  | *F. tardichlamydosporum* | c2 | n.d. | n.d. | + | - | + | *mat1-2* |
| cub0178 | Art3Art3 |  | Artemisa | Manzano vietnamita | Pisang Awak |  | *F. tardichlamydosporum* | c2 | n.d. | n.d. | + | - | + | *mat1-2* |
| cub0179 | ManVC3 |  | Villa Clara | Manzano vietnamita | Pisang Awak |  | *F. tardichlamydosporum* | c2 | n.d. | n.d. | + | - | + | *mat1-2* |
| cub0180 | R-1 |  | Villa Clara | Manzano | Silk |  | *F. tardichlamydosporum* | c2 | 0124 | R1 | + | - | + | *mat1-2* |

^1^- Code in the microbial culture collection of the Instituto de Investigaciones de Sanidad Vegetal (INISAV, Cuba);

^2,3^- According to DArT analysis;

^4,5^- According to Battle and Pérez-Vicente (2009);

*-determined in the present study;

Fox^a^-results according to the protocol described by Edel et al. (2000) with primers PFO2/PFO3;

TR4^b^- results of the PCR reaction with primers FocTR4-F/FocTR4-R and TR4^c^- with primers Six1a_266-F/ Six1a_266-R (Dita et al., 2010; Carvalhais et al., 2019);

R1/R2^d^-identified with the protocol described by (Carvalhais et al., 2019) and with primers Six6b_210-F/Six6b_210-R.
