## Supplementary material for "A deep genetic analysis of banana Fusarium wilt pathogens of Cuba in a Latin American and Caribbean diversity landscape": Table S4

**Table S4**. Origin details of a global collection of 243 *Fusarium* strains, that cause Fusarium wilt of banana, included in DArT analysis. Their proposed *Fusarium* species name derived from genotyping-by-sequencing (DArTseq) data and attributed clade, vegetative compatibility group, and race identification.

| **Code** | |  | **Host** | | | **Species** | **VCG^1^** | **Clade** | **Race^2^** |
| --- | --- | --- | --- | --- | --- | --- | --- | --- | --- |
| **DArT id** | **Alternative Code** | **Country** | **variety** | **Subgroup** | **Genotype** |  |  |  |  |
| bra0001 | BR13 | Brazil | n.d. | n.d. | n.d. | *F. phialophorum* | 0120/15 | c1 | n.d. |
| bra0003 | Pacovan | Brazil | Pacovan | Silk | AAB | *F. phialophorum* | 0120 | c1 | n.d. |
| bra0002 | GMB | Brazil | Gros Michel | Gros Michel | AAA | *F. tardichlamydosporum* | 0124 | c2 | n.d. |
| bra0004 | FocCNPMF-R1 | Brazil | Silk | Silk | AAB | *F. oxysporum* | n.d. | c3 | R1 |
| bra0005 | FocCNPMF-R2 | Brazil | Monthan | Monthan | ABB | *F. tardichlamydosporum* | 0124 | c2 | R2 |
| col0001 | 1 | Colombia | Gros Michel | Gros Michel | AAA | *F. phialophorum* | n.d. | c1 | n.d. |
| col0002 | 2 | Colombia | Platano Hawaiano | Cavendish | AAA | *F. phialophorum* | n.d. | c1 | n.d. |
| col0003 | 3_1 | Colombia | Platano Hawaiano | Cavendish | AAA | *F. phialophorum* | n.d. | c1 | n.d. |
| col0004 | 4_1 | Colombia | Gros Michel | Gros Michel | AAA | *F. phialophorum* | n.d. | c1 | n.d. |
| col0005 | 5 | Colombia | Gros Michel | Gros Michel | AAA | *F. oxysporum* | n.d. | c3 | n.d. |
| col0006 | 6_1 | Colombia | Gros Michel | Gros Michel | AAA | *F. phialophorum* | n.d. | c1 | n.d. |
| col0007 | 7 | Colombia | Gros Michel | Gros Michel | AAA | *F. phialophorum* | n.d. | c1 | n.d. |
| col0008 | 8_1 | Colombia | Gros Michel | Gros Michel | AAA | *F. phialophorum* | n.d. | c1 | n.d. |
| col0009 | 9_1 | Colombia | Gros Michel | Gros Michel | AAA | *F. phialophorum* | n.d. | c1 | n.d. |
| col0012 | 12 | Colombia | Platano Harton | n.d. | n.d. | *F. oxysporum* | n.d. | c3 | n.d. |
| col0013 | 13_1 | Colombia | Popocho | Bluggoe | ABB | *F. tardichlamydosporum* | n.d. | c2 | n.d. |
| col0014 | 14_1 | Colombia | Gros Michel | Gros Michel | AAA | *F. tardichlamydosporum* | n.d. | c2 | n.d. |
| col0015 | 15 | Colombia | Banano Urabeno | Cavendish | AAA | *F. tardicrescens* | n.d. | c2 | n.d. |
| col0016 | 16 | Colombia | Banano Urabeno | Cavendish | AAA | *F. phialophorum* | n.d. | c1 | n.d. |
| col0017 | 17 | Colombia | Banano Comun | Gros Michel | AAA | *F. phialophorum* | n.d. | c1 | n.d. |
| col0018 | 18 | Colombia | Banano Urabeno | Cavendish | AAA | *F. tardicrescens* | n.d. | c2 | n.d. |
| col0019 | 19_1 | Colombia | Guineo Mampora | Cavendish | AAA | *F. phialophorum* | n.d. | c1 | n.d. |
| col0020 | 20 | Colombia | Gros Michel | Gros Michel | AAA | *F. phialophorum* | n.d. | c1 | n.d. |
| col0021 | 21 | Colombia | Gros Michel | Gros Michel | AAA | *F. phialophorum* | n.d. | c1 | n.d. |

**Table S3**. (Continued).

| **Code** | |  | **Host** | | | **Species** | **VCG^1^** | **Clade** | **Race^2^** |
| --- | --- | --- | --- | --- | --- | --- | --- | --- | --- |
| **DArT id** | **Alternative Code** | **Country** | **variety** | **Subgroup** | **Genotype** |  |  |  |  |
| col0022 | 22 | Colombia | Gros Michel | Gros Michel | AAA | *F. phialophorum* | n.d. | c1 | n.d. |
| col0023 | 23_1 | Colombia | Gros Michel | Gros Michel | AAA | *F. phialophorum* | n.d. | c1 | n.d. |
| col0024 | 24_1 | Colombia | Banano Urabeno | Cavendish | AAA | *F. phialophorum* | n.d. | c1 | n.d. |
| col0025 | 25 | Colombia | Gros Michel | Gros Michel | AAA | *F. phialophorum* | n.d. | c1 | n.d. |
| col0026 | 26 | Colombia | Gros Michel | Gros Michel | AAA | *F. phialophorum* | n.d. | c1 | n.d. |
| col0027 | 27 | Colombia | Gros Michel | Gros Michel | AAA | *F. phialophorum* | n.d. | c1 | n.d. |
| col0028 | 28 | Colombia | Gros Michel | Gros Michel | AAA | *F. grosmichelii* | n.d. | c2 | n.d. |
| col0029 | 29 | Colombia | Ban.d.no Comun | Gros Michel | AAA | *F. phialophorum* | n.d. | c1 | n.d. |
| col0030 | 30 | Colombia | Gros Michel | Gros Michel | AAA | *F. phialophorum* | n.d. | c1 | n.d. |
| col0031 | 31_1 | Colombia | Gros Michel | Gros Michel | AAA | *F. phialophorum* | n.d. | c1 | n.d. |
| col0032 | 32_1 | Colombia | n.d. | n.d. | NA | *F. tardicrescens* | n.d. | c2 | n.d. |
| col0033 | 33_1 | Colombia | Gros Michel | Gros Michel | AAA | *F. phialophorum* | n.d. | c1 | n.d. |
| col0034 | 34_1 | Colombia | Banano Criollo | Gros Michel | AAA | *F. phialophorum* | n.d. | c1 | n.d. |
| col0035 | 35_1 | Colombia | Banano Criollo | Gros Michel | AAA | *F. phialophorum* | n.d. | c1 | n.d. |
| col0036 | 36 | Colombia | n.d. | n.d. | NA | *F. grosmichelii* | n.d. | c2 | n.d. |
| col0037 | 37 | Colombia | n.d. | n.d. | NA | *F. phialophorum* | n.d. | c1 | n.d. |
| col0038 | 38 | Colombia | Manzano | Silk | AAB | *F. oxysporum* | n.d. | c3 | n.d. |
| col0039 | 39 | Colombia | Manzano | Silk | AAB | *F. tardicrescens* | n.d. | c2 | n.d. |
| col0040 | 40 | Colombia | Manzano | Silk | AAB | *F. oxysporum* | n.d. | c3 | n.d. |
| col0041 | 41 | Colombia | Manzano | Silk | AAB | *F. grosmichelii* | n.d. | c2 | n.d. |
| col0042 | 42 | Colombia | Manzano | Silk | AAB | *F. tardicrescens* | n.d. | c2 | n.d. |
| col0043 | 43 | Colombia | Manzano | Silk | AAB | *F. grosmichelii* | n.d. | c2 | n.d. |
| col0044 | 44_1 | Colombia | Manzano | Silk | AAB | *F. tardicrescens* | n.d. | c2 | n.d. |
| col0045 | 45 | Colombia | Manzano | Silk | AAB | *F. tardicrescens* | n.d. | c2 | n.d. |
| col0046 | 46 | Colombia | Manzano | Silk | AAB | *F. tardicrescens* | n.d. | c2 | n.d. |
| col0047 | 47 | Colombia | Manzano | Silk | AAB | *F. tardicrescens* | n.d. | c2 | n.d. |

**Table S3**. (Continued).

| **Code** | |  | **Host** | | | | **Species** | **VCG^1^** | **Clade** | **Race^2^** |
| --- | --- | --- | --- | --- | --- | --- | --- | --- | --- | --- |
| **DArT id** | **Alternative Code** | **Country** | **variety** | **Subgroup** | **Genotype** | |  |  |  |  |
| col0048 | 48 | Colombia | Manzano | Silk | | AAB | *F. tardicrescens* | n.d. | c2 | n.d. |
| col0050 | 50_1 | Colombia | Manzano | Silk | | AAB | *F. tardicrescens* | n.d. | c2 | n.d. |
| col0051 | 51 | Colombia | Manzano | Silk | | AAB | *F. tardicrescens* | n.d. | c2 | n.d. |
| col0052 | 52_1 | Colombia | Manzano | Silk | | AAB | *F. tardicrescens* | n.d. | c2 | n.d. |
| col0053 | 53 | Colombia | Manzano | Silk | | AAB | *F. tardicrescens* | n.d. | c2 | n.d. |
| col0054 | 54 | Colombia | Manzano | Silk | | AAB | *F. tardicrescens* | n.d. | c2 | n.d. |
| col0055 | 55 | Colombia | Popocho | Bluggoe | | ABB | *F. tardichlamydosporum* | n.d. | c2 | n.d. |
| col0056 | 56 | Colombia | Manzano | Silk | | AAB | *F. oxysporum* | n.d. | c1 | n.d. |
| col0057 | 57 | Colombia | Manzano | Silk | | AAB | *F. phialophorum* | n.d. | c1 | n.d. |
| col0059 | 59_1 | Colombia | Manzano | Silk | | AAB | *F. tardicrescens* | n.d. | c2 | n.d. |
| col0060 | 60 | Colombia | Manzano | Silk | | AAB | *F. tardicrescens* | n.d. | c2 | n.d. |
| col0061 | 61 | Colombia | Gros Michel | Gros Michel | | AAA | *F. phialophorum* | n.d. | c1 | n.d. |
| col0063 | 63 | Colombia | Gros Michel | Gros Michel | | AAA | *F. phialophorum* | n.d. | c1 | n.d. |
| col0065 | 65 | Colombia | Gros Michel | Gros Michel | | AAA | *F. phialophorum* | n.d. | c1 | n.d. |
| col0066 | 66 | Colombia | Gros Michel | Gros Michel | | AAA | *F. phialophorum* | n.d. | c1 | n.d. |
| col0067 | 67 | Colombia | Gros Michel | Gros Michel | | AAA | *F. tardicrescens* | n.d. | c2 | n.d. |
| col0069 | 69 | Colombia | Gros Michel | Gros Michel | | AAA | *F. phialophorum* | n.d. | c1 | n.d. |
| col0070 | 70 | Colombia | Gros Michel | Gros Michel | | AAA | *F. phialophorum* | n.d. | c1 | n.d. |
| col0071 | 71 | Colombia | Gros Michel | Gros Michel | | AAA | *F. phialophorum* | n.d. | c1 | n.d. |
| col0072 | 72 | Colombia | Gros Michel | Gros Michel | | AAA | *F. phialophorum* | n.d. | c1 | n.d. |
| col0073 | 73 | Colombia | Gros Michel | Gros Michel | | AAA | *F. phialophorum* | n.d. | c1 | n.d. |
| col0074 | 74 | Colombia | n.d. | n.d. | | NA | *F. tardicrescens* | n.d. | c2 | n.d. |
| col0075 | 75 | Colombia | Gros Michel | Gros Michel | | AAA | *F. phialophorum* | n.d. | c1 | n.d. |
| col0077 | 77 | Colombia | n.d. | n.d. | | NA | *F. phialophorum* | n.d. | c1 | n.d. |
| col0082 | 82 | Colombia | Manzano | Silk | | AAB | *F. tardicrescens* | n.d. | c2 | n.d. |
| col0083 | 83 | Colombia | Manzano | Silk | | AAB | *F. tardicrescens* | n.d. | c2 | n.d. |

**Table S3**. (Continued).

| **Code** | |  | **Host** | | | | **Species** | **VCG^1^** | **Clade** | **Race^2^** |
| --- | --- | --- | --- | --- | --- | --- | --- | --- | --- | --- |
| **DArT id** | **Alternative Code** | **Country** | **variety** | **Subgroup** | **Genotype** | |  |  |  |  |
| cos0004 | 103 | Costa Rica | Gros Michel | Gros Michel | | AAA | *F. phialophorum* | 0120/15 | c1 | n.d. |
| cos0049 | 56_1 | Costa Rica | Gros Michel | Gros Michel | | AAA | *F. phialophorum* | 0120/15 | c1 | n.d. |
| cos0052 | 59_2 | Costa Rica | Gros Michel | Gros Michel | | AAA | *F. phialophorum* | 0120/15 | c1 | n.d. |
| cos0063 | 70_1 | Costa Rica | Gros Michel | Gros Michel | | AAA | *F. phialophorum* | 0120/15 | c1 | n.d. |
| cos0070 | 77_1 | Costa Rica | Gros Michel | Gros Michel | | AAA | *F. phialophorum* | 0120/15 | c1 | n.d. |
| cos0084 | 92 | Costa Rica | Gros Michel | Gros Michel | | AAA | *F. phialophorum* | 0120/15 | c1 | n.d. |
| cos0092 | BIF-01 | Costa Rica | n.d. | n.d. | | NA | *F. phialophorum* | 0120/15 | c1 | n.d. |
| cos0094 | BIF-04 | Costa Rica | n.d. | n.d. | | NA | *F. phialophorum* | 0120/15 | c1 | n.d. |
| cos0095 | BIF-05 | Costa Rica | n.d. | n.d. | | NA | *F. phialophorum* | newVCG1 | c1 | n.d. |
| cos0097 | BIF-11 | Costa Rica | n.d. | n.d. | | NA | *F. phialophorum* | 0120/15 | c1 | n.d. |
| cos0098 | BIF-16 | Costa Rica | n.d. | n.d. | | NA | *F. phialophorum* | 0120/15 | c1 | n.d. |
| cos0099 | BIF-19 | Costa Rica | n.d. | n.d. | | NA | *F. phialophorum* | 0120/15 | c1 | n.d. |
| cos0100 | BIF-21 | Costa Rica | n.d. | n.d. | | NA | *F. phialophorum* | 0120/15 | c1 | n.d. |
| cos0101 | BIF-24 | Costa Rica | n.d. | n.d. | | NA | *F. phialophorum* | 0120/15 | c1 | n.d. |
| cos0102 | BIF-27 | Costa Rica | n.d. | n.d. | | NA | *F. phialophorum* | 0120/15 | c1 | n.d. |
| cos0103 | BIF-28 | Costa Rica | n.d. | n.d. | | NA | *F. phialophorum* | 0120/15 | c1 | n.d. |
| cos0104 | BIF-29 | Costa Rica | n.d. | n.d. | | NA | *F. phialophorum* | 0120/15 | c1 | n.d. |
| cos0105 | BIF-30 | Costa Rica | n.d. | n.d. | | NA | *F. phialophorum* | 0120/15 | c1 | n.d. |
| cos0106 | BIF-38 | Costa Rica | n.d. | n.d. | | NA | *F. phialophorum* | 0120/15 | c1 | n.d. |
| cos0107 | BIF-41 | Costa Rica | n.d. | n.d. | | NA | *F. phialophorum* | 0120/15 | c1 | n.d. |
| cos0108 | BIF-43 | Costa Rica | n.d. | n.d. | | NA | *F. phialophorum* | 0120/15 | c1 | n.d. |
| cos0109 | BIF-44 | Costa Rica | n.d. | n.d. | | NA | *F. phialophorum* | 0120/15 | c1 | n.d. |
| cos0110 | BIF-54 | Costa Rica | n.d. | n.d. | | NA | *F. phialophorum* | 0120/15 | c1 | n.d. |
| cos0111 | BIF-55 | Costa Rica | n.d. | n.d. | | NA | *F. phialophorum* | 0120/15 | c1 | n.d. |
| cos0112 | BIF-58 | Costa Rica | n.d. | n.d. | | NA | *F. phialophorum* | 0120/15 | c1 | n.d. |
| cos0113 | BIF-60 | Costa Rica | n.d. | n.d. | | NA | *F. phialophorum* | 0120/15 | c1 | n.d. |

**Table S3**. (Continued).

| **Code** | |  | **Host** | | | | **Species** | **VCG^1^** | **Clade** | **Race^2^** |
| --- | --- | --- | --- | --- | --- | --- | --- | --- | --- | --- |
| **DArT id** | **Alternative Code** | **Country** | **variety** | **Subgroup** | **Genotype** | |  |  |  |  |
| cos0114 | BIF-77 | Costa Rica | n.d. | n.d. | | NA | *F. tardicrescens* | newVCG2 | c2 | n.d. |
| cos0115 | BIF-78 | Costa Rica | Gros Michel | Gros Michel | | AAA | *F. phialophorum* | 0120/15 | c1 | n.d. |
| cos0116 | CR1.1A | Costa Rica | Gros Michel | Gros Michel | | AAA | *F. phialophorum* | 0120 | c1 | R1 |
| cos0117 | Foc1 | Costa Rica | Gros Michel | Gros Michel | | AAA | *F. phialophorum* | 0120/15 | c1 | n.d. |
| cos0118 | Foc14 | Costa Rica | Gros Michel | Gros Michel | | AAA | *F. phialophorum* | 0120/15 | c1 | n.d. |
| cos0119 | Foc19508 | Costa Rica | Gros Michel | Gros Michel | | AAA | *F. phialophorum* | 0120 | c1 | n.d. |
| cos0120 | FocD3 | Costa Rica | n.d. | n.d. | | NA | *F. phialophorum* | 0120/15 | c1 | n.d. |
| cos0121 | FocD3 | Costa Rica | n.d. | n.d. | | NA | *F. phialophorum* | 0120/15 | c1 | n.d. |
| cos0122 | FD8 | Costa Rica | n.d. | n.d. | | NA | *F. phialophorum* | 0120/15 | c1 | n.d. |
| cos0124 | FocP1 | Costa Rica | n.d. | n.d. | | NA | *F. phialophorum* | 0120/15 | c1 | n.d. |
| cos0125 | FocP11 | Costa Rica | n.d. | n.d. | | NA | *F. phialophorum* | 0120/15 | c1 | n.d. |
| cos0126 | FocP12 | Costa Rica | n.d. | n.d. | | NA | *F. phialophorum* | 0120/15 | c1 | n.d. |
| cos0127 | FocP13 | Costa Rica | n.d. | n.d. | | NA | *F. phialophorum* | 0120/15 | c1 | n.d. |
| cos0128 | FocP17 | Costa Rica | n.d. | n.d. | | NA | *F. phialophorum* | 0120/15 | c1 | n.d. |
| cos0130 | FocP7 | Costa Rica | n.d. | n.d. | | NA | *F. phialophorum* | 0120/15 | c1 | n.d. |
| cos0131 | Focu2 | Costa Rica | Gros Michel | Gros Michel | | AAA | *F. phialophorum* | 0120 | c1 | n.d. |
| cos0132 | FS2 | Costa Rica | n.d. | n.d. | | NA | *F. phialophorum* | 0120/15 | c1 | n.d. |
| cos0133 | STGM1 | Costa Rica | Gros Michel | Gros Michel | | AAA | *F. phialophorum* | 0120 | c1 | R1 |
| hon0001 | 34661 | Honduras | Highgate | Gros Michel | | AAA | *F. phialophorum* | 0120 | c1 | R1 |
| hon0003 | NRRL36107 | Honduras | Maqueño | Plantain | | AAB | *F. purpurascens* | 0126 | c1 | ST4 |
| hon0005 | S1 | Honduras | Highgate | Gros Michel | | AAA | *F. purpurascens* | 0126 | c1 | n.d. |
| hon0002 | NRRL36105 | Honduras | n.d. | Bluggoe | | ABB | *F. tardichlamydosporum* | 0124 | c2 | R2 |
| hon0004 | Focu3 | Honduras | n.d. | Bluggoe | | ABB | *F. tardichlamydosporum* | 0124 | c2 | n.d. |
| hon0006 | STD1 | Honduras | Highgate | Gros Michel | | AAA | *F. tardichlamydosporum* | 0124 | c2 | R1 |
| jam0003 | FCJ7 | Jamaica | n.d. | Cavendish | | AAA | *F. phialophorum* | 0120 | c1 | n.d. |
| jam0001 | 1S.92 | Jamaica | Williams | Cavendish | | AAA | *F. tardichlamydosporum* | 0125 | c2 | n.d. |

**Table S3**. (Continued).

| **Code** | |  | **Host** | | | | **Species** | **VCG^1^** | **Clade** | **Race^2^** |
| --- | --- | --- | --- | --- | --- | --- | --- | --- | --- | --- |
| **DArT id** | **Alternative Code** | **Country** | **variety** | **Subgroup** | **Genotype** | |  |  |  |  |
| jam0002 | FCJ2 | Jamaica | n.d. | Bluggoe | | ABB | *F. tardichlamydosporum* | 0124 | c2 | n.d. |
| jam0004 | Focu4 | Jamaica | n.d. | Bluggoe | | ABB | *F. tardichlamydosporum* | 0124 | c2 | n.d. |
| mex0002 | Mex12 | Mexico | Gros Michel | Gros Michel | | AAA | *F. tardichlamydosporum* | 0124/5/8 | c2 | n.d. |
| mex0001 | Mex14 | Mexico | n.d. | Bluggoe | | ABB | *F. tardichlamydosporum* | 0124/5/8 | c2 | n.d. |
| nic0006 | Foc16 | Nicaragua | Gros Michel | Gros Michel | | AAA | *F. oxysporum* | newVCG13 | c3 | n.d. |
| nic0010 | Foc8 | Nicaragua | Gros Michel | Gros Michel | | AAA | *F. oxysporum* | newVCG12 | c3 | n.d. |
| nic0001 | BPnic0901 | Nicaragua | Cuadrado | Bluggoe | | ABB | *F. tardichlamydosporum* | 0124/5 | c2 | n.d. |
| nic0002 | BPNic0902 | Nicaragua | Cuadrado | Bluggoe | | ABB | *F. tardichlamydosporum* | 0124/5 | c2 | n.d. |
| nic0003 | BPNic0903 | Nicaragua | Cuadrado | Bluggoe | | ABB | *F. tardichlamydosporum* | 0124/5 | c2 | n.d. |
| nic0011 | Nic10.01 | Nicaragua | n.d. | Bluggoe | | ABB | *F. tardichlamydosporum* | 0124/5 | c2 | n.d. |
| nic0012 | Nic10.02 | Nicaragua | n.d. | Bluggoe | | ABB | *F. tardichlamydosporum* | 0124/5 | c2 | n.d. |
| nic0013 | Nic10.04 | Nicaragua | n.d. | Bluggoe | | ABB | *F. tardichlamydosporum* | 0124/5 | c2 | n.d. |
| nic0014 | Nic11 | Nicaragua | n.d. | Bluggoe | | ABB | *F. tardichlamydosporum* | 0124/5/8 | c2 | n.d. |
| nic0015 | STN1 | Nicaragua | n.d. | Bluggoe | | ABB | *F. tardichlamydosporum* | 0124 | c2 | n.d. |
| per0001 | P104 | Peru | Isla | Iholena | | AAB | *F. tardicrescens* | newVCG8 | c2 | n.d. |
| per0002 | P107 | Peru | Isla | Iholena | | AAB | *F. phialophorum* | 0120/15 | c1 | n.d. |
| per0003 | P113b | Peru | Manzano | Silk | | AAB | *F. oxysporum* | newVCG16 | c3 | n.d. |
| per0004 | P113c | Peru | Manzano | Silk | | AAB | *F. tardicrescens* | newVCG8 | c2 | n.d. |
| per0005 | P113d | Peru | Manzano | Silk | | AAB | *F. phialophorum* | 0120/15 | c1 | n.d. |
| per0006 | P114 | Peru | Manzano | Silk | | AAB | *F. phialophorum* | 0120/15 | c1 | n.d. |
| per0007 | P131 | Peru | Morado | Red | | AAA | *F. phialophorum* | 0120/15 | c1 | n.d. |
| per0008 | P132b | Peru | Manzano | Silk | | AAB | *F. tardicrescens* | newVCG8 | c2 | n.d. |
| per0009 | P132d | Peru | Manzano | Silk | | AAB | *F. oxysporum* | newVCG17 | c3 | n.d. |
| per0010 | P133 | Peru | Morado | Red | | AAA | *F. phialophorum* | 0120/15 | c1 | n.d. |
| per0011 | P135 | Peru | Palillo | Maoli-Popoulu | | AAB | *F. oxysporum* | newVCG18 | c3 | n.d. |
| per0015 | P14 | Peru | Isla | Iholena | | AAB | *F. tardicrescens* | newVCG7 | c2 | n.d. |

**Table S3**. (Continued).

| **Code** | |  | **Host** | | | | **Species** | **VCG^1^** | **Clade** | **Race^2^** |
| --- | --- | --- | --- | --- | --- | --- | --- | --- | --- | --- |
| **DArT id** | **Alternative Code** | **Country** | **variety** | **Subgroup** | **Genotype** | |  |  |  |  |
| per0016 | P15a | Peru | Isla | Iholena | | AAB | *F. oxysporum* | newVCG17 | c3 | n.d. |
| per0017 | P15d | Peru | Isla | Iholena | | AAB | *F. oxysporum* | newVCG17 | c3 | n.d. |
| per0018 | P15d_1 | Peru | Isla | Iholena | | AAB | *F. phialophorum* | newVCG17 | c1 | n.d. |
| per0019 | P17 | Peru | Isla | Iholena | | AAB | *F. tardicrescens* | newVCG7 | c2 | n.d. |
| per0020 | P18 | Peru | Manzano | Silk | | AAB | *F. tardicrescens* | newVCG7 | c2 | n.d. |
| per0021 | P20a | Peru | Manzano | Silk | | AAB | *F. oxysporum* | newVCG16 | c3 | n.d. |
| per0022 | P20d | Peru | Manzano | Silk | | AAB | *F. tardicrescens* | newVCG8 | c2 | n.d. |
| per0023 | P21 | Peru | Manzano | Silk | | AAB | *F. oxysporum* | newVCG16 | c3 | n.d. |
| per0024 | P22 | Peru | Manzano | Silk | | AAB | *F. tardicrescens* | newVCG7 | c2 | n.d. |
| per0025 | P24a | Peru | Seda | Gros Michel | | AAA | *F. tardicrescens* | newVCG7 | c2 | n.d. |
| per0026 | P24c | Peru | Seda | Gros Michel | | AAA | *F. phialophorum* | 0120/15 | c1 | n.d. |
| per0027 | P25 | Peru | Seda | Gros Michel | | AAA | *F. oxysporum* | newVCG18 | c3 | n.d. |
| per0029 | P26 | Peru | Seda | Gros Michel | | AAA | *F. oxysporum* | newVCG17 | c3 | n.d. |
| per0030 | P29a | Peru | Isla | Iholena | | AAB | *F. phialophorum* | 0120/15 | c1 | n.d. |
| per0031 | P29b | Peru | Isla | Iholena | | AAB | *F. oxysporum* | newVCG16 | c3 | n.d. |
| per0032 | P2c | Peru | Isla | Iholena | | AAB | *F. phialophorum* | 0120/15 | c1 | n.d. |
| per0033 | P2d | Peru | Isla | Iholena | | AAB | *F. oxysporum* | newVCG16 | c3 | n.d. |
| per0034 | P30 | Peru | Isla | Iholena | | AAB | *F. phialophorum* | 0120/15 | c1 | n.d. |
| per0035 | P34 | Peru | Isla | Iholena | | AAB | *F. phialophorum* | 0120/15 | c1 | n.d. |
| per0036 | P37 | Peru | Isla | Iholena | | AAB | *F. tardicrescens* | newVCG8 | c2 | n.d. |
| per0037 | P38 | Peru | Isla | Iholena | | AAB | *F. phialophorum* | 0120/15 | c1 | n.d. |
| per0038 | P38_1 | Peru | Isla | Iholena | | AAB | *F. phialophorum* | 0120/15 | c1 | n.d. |
| per0039 | P3 | Peru | Isla | Iholena | | AAB | *F. phialophorum* | 0120/15 | c1 | n.d. |
| per0040 | P41 | Peru | Seda | Gros Michel | | AAA | *F. tardicrescens* | newVCG8 | c2 | n.d. |
| per0041 | P47 | Peru | Isla | Iholena | | AAB | *F. tardichlamydosporum* | 0124/5 | c2 | n.d. |
| per0042 | P47a | Peru | Isla | Iholena | | AAB | *F. phialophorum* | 0120/15 | c1 | n.d. |

**Table S3**. (Continued).

| **Code** | |  | **Host** | | | | **Species** | **VCG^1^** | **Clade** | **Race^2^** |
| --- | --- | --- | --- | --- | --- | --- | --- | --- | --- | --- |
| **DArT id** | **Alternative Code** | **Country** | **variety** | **Subgroup** | **Genotype** | |  |  |  |  |
| per0043 | P49 | Peru | Isla | Iholena | | AAB | *F. tardicrescens* | newVCG7 | c2 | n.d. |
| per0044 | P50 | Peru | Isla | Iholena | | AAB | *F. tardicrescens* | newVCG7 | c2 | n.d. |
| per0045 | per0045 | Peru | Seda | Gros Michel | | AAA | *F. tardicrescens* | newVCG8 | c2 | n.d. |
| per0046 | per0046 | Peru | Seda | Gros Michel | | AAA | *F. oxysporum* | newVCG17 | c3 | n.d. |
| per0047 | per0047 | Peru | Isla | Iholena | | AAB | *F. phialophorum* | 0120/15 | c1 | n.d. |
| per0048 | per0048 | Peru | Isla | Iholena | | AAB | *F. phialophorum* | 0120/15 | c1 | n.d. |
| per0049 | per0049 | Peru | Isla | Iholena | | AAB | *F. phialophorum* | 0120/15 | c1 | n.d. |
| per0050 | per0050 | Peru | Isla | Iholena | | AAB | *F. tardicrescens* | newVCG7 | c2 | n.d. |
| per0051 | per0051 | Peru | Isla | Iholena | | AAB | *F. oxysporum* | newVCG18 | c3 | n.d. |
| per0052 | per0052 | Peru | Isla | Iholena | | AAB | *F. oxysporum* | newVCG17 | c3 | n.d. |
| per0053 | per0053 | Peru | Isla | Iholena | | AAB | *F. tardicrescens* | newVCG7 | c2 | n.d. |
| per0054 | per0054 | Peru | Morado | Red | | AAA | *F. phialophorum* | 0120/15 | c1 | n.d. |
| per0055 | per0055 | Peru | Morado | Red | | AAA | *F. tardicrescens* | newVCG7 | c2 | n.d. |
| per0056 | per0056 | Peru | Seda | Gros Michel | | AAA | *F. tardicrescens* | newVCG7 | c2 | n.d. |
| per0057 | per0057 | Peru | Isla | Iholena | | AAB | *F. oxysporum* | newVCG17 | c3 | n.d. |
| per0058 | per0058 | Peru | Seda | Gros Michel | | AAA | *F. phialophorum* | 0120/15 | c1 | n.d. |
| per0059 | per0059 | Peru | Isla | Iholena | | AAB | *F. phialophorum* | 0120/15 | c1 | n.d. |
| per0060 | per0060 | Peru | Isla | Iholena | | AAB | *F. oxysporum* | newVCG17 | c3 | n.d. |
| per0061 | per0061 | Peru | Isla | Iholena | | AAB | *F. oxysporum* | newVCG19 | c3 | n.d. |
| per0062 | per0062 | Peru | Isla | Iholena | | AAB | *F. tardicrescens* | newVCG9 | c2 | n.d. |
| per0063 | per0063 | Peru | Isla | Iholena | | AAB | *F. tardicrescens* | newVCG7 | c2 | n.d. |
| per0064 | per0064 | Peru | Isla | Iholena | | AAB | *F. tardicrescens* | newVCG7 | c2 | n.d. |
| per0065 | per0065 | Peru | Isla | Iholena | | AAB | *F. tardicrescens* | newVCG7 | c2 | n.d. |
| usa0001 | A2-1 | USA | Apple | Silk | | AAB | *F. purpurascens* | 01210 | c1 | n.d. |
| usa0005 | CVA | USA | Apple | Silk | | AAB | *F. tardichlamydosporum* | 0124/5 | c2 | n.d. |
| usa0006 | Focu7 | USA | Apple | Silk | | AAB | *F. purpurascens* | 01210 | c1 | n.d. |

**Table S3**. (Continued).

| **Code** | |  | **Host** | | | | **Species** | **VCG^1^** | **Clade** | **Race^2^** |
| --- | --- | --- | --- | --- | --- | --- | --- | --- | --- | --- |
| **DArT id** | **Alternative Code** | **Country** | **variety** | **Subgroup** | **Genotype** | |  |  |  |  |
| usa0007 | IPO98-05 | USA | Bluggoe | Bluggoe | | ABB | *F. tardichlamydosporum* | 0124 | c2 | n.d. |
| usa0008 | MUCL38369 | USA | n.d. | n.d. | | NA | *F. tardichlamydosporum* | 0124/5 | c2 | n.d. |
| usa0009 | MUCL38370 | USA | n.d. | n.d. | | NA | *F. tardichlamydosporum* | 0124/5 | c2 | n.d. |
| usa0010 | PLBL | USA | n.d. | Bluggoe | | ABB | *F. tardichlamydosporum* | 0124/5/8 | c2 | n.d. |
| aus0018 | 24662 | Australia | n.d. | Cavendish | | AAA | *F. odoratissimum* | 01213 | c1 | TR4 |
| aus0019 | 24663 | Australia | n.d. | Cavendish | | AAA | *F. odoratissimum* | 01213 | c1 | TR4 |
| aus0023 | NRRL36109 | Australia | SH-3142 | FHIA hybrid | | AA | *F. phialophorum* | 01211 | c1 | ST4 |
| aus0024 | NRRL36110 | Australia | Mons Mari | Cavendish | | AAA | *F. purpurascens* | 0129 | c1 | ST4 |
| aus0032 | Focu1 | Australia | Mons Mari | Cavendish | | AAA | *F. phialophorum* | 0120 | c1 | n.d. |
| chi0001 | NRRL36102 | China | n.d. | Cavendish | | AAA | *F. odoratissimum* | 0121 | c1 | TR4 |
| ido0017 | InaCC F917 | Indonesia | Pisang Ambon | Cavendish | | AAA | *F. kalimantanense* | n.d. | c4 | n.p. |
| ido0018 | InaCC F918 | Indonesia | Pisang Ambon | Cavendish | | AAA | *F. kalimantanense* | n.d. | c4 | n.d. |
| ido0022 | InaCC F922 | Indonesia | Pisang Ambon | Cavendish | | AAA | *F. kalimantanense* | n.d. | c4 | n.d. |
| ido0062 | InaCC F960 | Indonesia | Pisang Kepok | Pisang awak | | ABB | *F. sangayamense* | n.d. | c4 | n.p. |
| ido0063 | InaCC F961 | Indonesia | Pisang Kepok | Pisang awak | | ABB | *F. sangayamense* | n.d. | c4 | n.d. |
| ido0089 | InaCC F817 | Indonesia | Pisang Kepok | Pisang awak | | ABB | *F. odoratissimum* | 01213 | c1 | TR4 |
| ido0095 | InaCC F984 | Indonesia | Pisang Kepok | Pisang awak | | ABB | *F. cugenangense* | n.d. | c2 | n.d. |
| ido0165 | InaCC F866 | Indonesia | Pisang Ambon Kuning | Gros Michel | | AAA | *F. hexaseptatum* | n.d. | c2 | R1 |
| ido0226 | II5 | Indonesia | Pisang Manurung | Pisang awak | | AAB | *F. odoratissimum* | 01213 | c1 | TR4 |
| ido0229 | Indo25 | Indonesia | Pisang Ambon | Cavendish | | AAA | *F. purpurascens* | 01219 | c1 | n.p. |
| ido0233 | InaCC F892 | Indonesia | Pisang Barangan | Lakatan | | AAA | *F. odoratissimum* | 01213/16 | c1 | TR4 |
| ido0234 | InaCC F904 | Indonesia | Pisang Kepok | Pisang awak | | AAA | *F. odoratissimum* | 01213 | c1 | TR4 |
| mas0002 | NRRL36115 | Malaysia | Pisang Ambon | Cavendish | | AAA | *F. duoseptatum* | 01224 | c2 | n.d. |
| mas0005 | Mal43 | Malaysia | Pisang Rastali | Silk | | AAB | *F. duoseptatum* | 01217 | c2 | R1 |
| mau0001 | Mau10-01 | Mauritius | Gingeli Bes | Silk | | ABB | *F. grosmichelii* | newVCG6 | c2 | n.d. |
| jor0001 | JV11 | Jordan | n.d. | Cavendish | | AAA | *F. odoratissimum* | 01213/16 | c1 | TR4 |

**Table S3**. (Continued).

| **Code** | |  | **Host** | | | | | **Species** | | **VCG^1^** | | **Clade** | | **Race^2^** |
| --- | --- | --- | --- | --- | --- | --- | --- | --- | --- | --- | --- | --- | --- | --- |
| **DArT id** | **Alternative Code** | **Country** | **variety** | **Subgroup** | **Genotype** | | |  |  |  |  |  |  |  |
| leb0001 | Leb1.2C | Lebanon | n.d. | Cavendish | | AAA | *F. odoratissimum* | | 01213 | | c1 | | TR4 | |
| mal0003 | Mal1.5b | Malawi | n.d. | n.d. | | NA | *F. oxysporum* | | newVCG15 | | c3 | | n.d. | |
| mal0001 | NRRL36113 | Malawi | Harare | Bluggoe | | ABB | *F. tardicrescens* | | 01214 | | c2 | | n.d. | |
| tha0001 | NRRL36118 | Thailand | Kluai nam wa | Pisang awak | | ABB | *F. cugenangense* | | 01221 | | c2 | | n.d. | |
| tha0002 | NRRL36120 | Thailand | Kluai nam wa | Pisang awak | | ABB | *F. grosmichelii* | | 01218 | | c2 | | R1 | |
| pak0001 | Pak1.1A | Pakistan | Cavendish | Cavendish | | AAA | *F. odoratissimum* | | 01213 | | c1 | | TR4 | |
| phi0002 | Phi1.1A | Philippines | Williams | Cavendish | | AAA | *F. odoratissimum* | | 01213 | | c1 | | TR4 | |
| phi0005 | Phi2.6C | Philippines | GCTCV-218 | Cavendish | | AAA | *F. odoratissimum* | | 01213 | | c1 | | TR4 | |
| phi0001 | NRRL36103 | Philippines | n.d. | Cavendish | | AAA | *F. purpurascens* | | 0122 | | c1 | | n.d. | |
| tan0001 | NRRL36108 | Tanzania | Ney Poovan | Ney Poovan | | AB | *F. tardichlamydosporum* | | 01212 | | c2 | | n.d. | |
| CBS221.76 | CBS221.76/Ff | NA | Rice | Non-Musa | | Rice | *F. fujikuroi* | | - | | - | | - | |
| Fol4287 | Fol4287 | NA | tomato | tomato | | Tomato | *F. oxysporum* | | - | | c3 | | - | |

^1^-VCG according to Ordoñez et al.(2018). ^2^-Race related (race 1, R1; race 2, R2; race 4, R4; subtropical race 4, ST4 and tropical race 4, TR4) to each isolate according to previous reports (Boehm et al., 1994; Fourie et al., 2009; Dita et al., 2010; Maryani et al., 2019). n.d. stands for "not determined" and n.p. for non to pathogenic to Grand Naine or Gros Michel.
